# Beech tree masting explains the inter-annual variation in the fall and spring peaks of *Ixodes ricinus* ticks with different time lags

**DOI:** 10.1101/2021.04.20.440690

**Authors:** Cindy Bregnard, Olivier Rais, Coralie Herrmann, Olaf Kahl, Katharina Brugger, Maarten J. Voordouw

## Abstract

**Background:** The tick *Ixodes ricinus* is an important vector of tick-borne diseases including Lyme borreliosis. In continental Europe, the nymphal stage of *I. ricinus* often has a bimodal phenology with a large spring/early summer peak and a smaller fall peak. While there is consensus about the origin of the spring nymphal peak, there are two alternative hypotheses for the fall nymphal peak, direct development versus delayed diapause. These two hypotheses make different predictions about the time lags of the correlations between the spring peak, the fall peak, and seed production (masting) by deciduous trees.

**Methods:** To determine which hypothesis is most important for explaining the fall peak, we used data from a long-term surveillance study (15 years) on the density of *I. ricinus* nymphal ticks at 4 different elevation sites in an area of Switzerland that is endemic for Lyme borreliosis, and long-term data on the mast of the European beech tree from the literature.

**Results:** *I. ricinus* nymphs had a bimodal phenology at the three lower elevation sites, but a unimodal phenology at the top elevation site. At the lower elevation sites, the density of nymphs (DON) in the fall was strongly correlated with the DON in the spring of the following year. The inter-annual variation in the densities of *I. ricinus* nymphs in the fall and spring were best explained by a 1-year versus a 2-year time lag with the beech tree masting index. Fall nymphs had higher fat content and are younger than spring nymphs. All of these observations are consistent with the direct development hypothesis for the fall peak of *I. ricinus* nymphs at our study site. Our study provides new insight into the complex bimodal phenology of this important disease vector.

**Conclusions:** Public health officials in Europe should be aware that following a strong mast year, the DON will increase 1 year later in the fall and 2 years later in the spring and summer. Population ecology studies of *I. ricinus* should consider that the spring and fall peak in the same calendar year represent different generations of ticks.

## INTRODUCTION

The incidence of tick-borne diseases is increasing in Europe and North America [1–7]. In large parts of Europe, the hard tick *Ixodes ricinus* is an important vector of a variety of tick-borne diseases including Lyme borreliosis and tick-borne encephalitis [8, 9]. This tick species consists of three motile stages, larva, nymph, and adult, that must obtain a blood meal from a vertebrate host to moult into the next stage (or produce eggs in the case of adult female ticks). The population ecology of *I. ricinus* is complicated by a number of factors. First, *I. ricinus* can feed on a wide variety of vertebrate hosts (e.g., lizards, rodents, birds, carnivores, ungulates), for which the population density is often unknown [10, 11]. Second, the life cycle takes several years to complete, which introduces time lags [12–14]. For example, the density of nymphs in year *y* depends on the feeding success of the larvae in year *y*-1, which depends on the ratio of larvae to vertebrate reservoir hosts in year *y*-1 [15, 16]. Third, the existence of diapause as an adaptation to surviving cold winters can split the same cohort of ticks into groups that are active at different times of the year [17–19]. Uncertainty about the origin of these groups complicates our ability to model the underlying ecological factors and appropriate time lags that drive inter-annual variation in tick abundance.

Long-term studies have shown that a combination of abiotic and biotic factors drive inter-annual variation in tick abundance. *Ixodes* ticks spend more than 99% of their life cycle off the host, where they are exposed to changes in temperature and precipitation [17, 20]. The life history traits of *Ixodes* ticks, such as development rates and survival rates, are highly sensitive to temperature and relative humidity (RH) [17, 21–23]. Tick population ecology is also highly sensitive to the abundance of vertebrate hosts because all tick stages must blood feed to graduate to the next stage in the life cycle [24, 25]. Small mammals (e.g., rodents) are an important but variable source of food for immature ticks (larvae and nymphs); rodent populations often exhibit inter-annual fluctuations due to variation in their food supply [26–31]. Studies on *I. ricinus* in Europe and on *I. scapularis* in North America have shown that inter-annual variation in seed production by deciduous trees drives inter-annual variation in the density of nymphs 2 years later; this relationship is mediated by rodents that feed on the tree seeds and provide blood meals for the larvae [12, 14, 15, 32–36]. In summary, masting in the fall of year *y*-2 enhances rodent density and larval feeding success in the spring and summer of year *y*-1, which increases the density of nymphs in year *y*.

*I. ricinus* has a distinct seasonal activity pattern (phenology) that allows them to search for vertebrate hosts (a behavior called questing) when abiotic conditions (temperature and humidity) are favorable. Diapause is a critical adaptation that allows ticks (and other arthropods) to overwinter in an inactive state and thereby avoid cold winter temperatures. Behavioral diapause is the suppression of host-seeking activity by unfed ticks in the fall in anticipation of unfavorable winter conditions. Developmental diapause is the cessation of development by engorged ticks in the fall to enhance overwinter survival [37]. Both types of diapause are driven by changes in photoperiod (the most reliable predictor of seasonal change) and both are important for structuring the phenology of *I. ricinus* ticks [19, 21], which varies widely among geographic locations [17-19, 38, 39]. In some parts of Europe, nymphs and adult ticks exhibit a unimodal phenology where questing activity peaks in late spring or early summer and ends in the fall (Table 1) [40–44]. In central Europe, the most common phenology is bimodal with a large peak of activity in spring/early summer and a smaller peak of activity in fall (Table 1) [19].

**Table 1.**
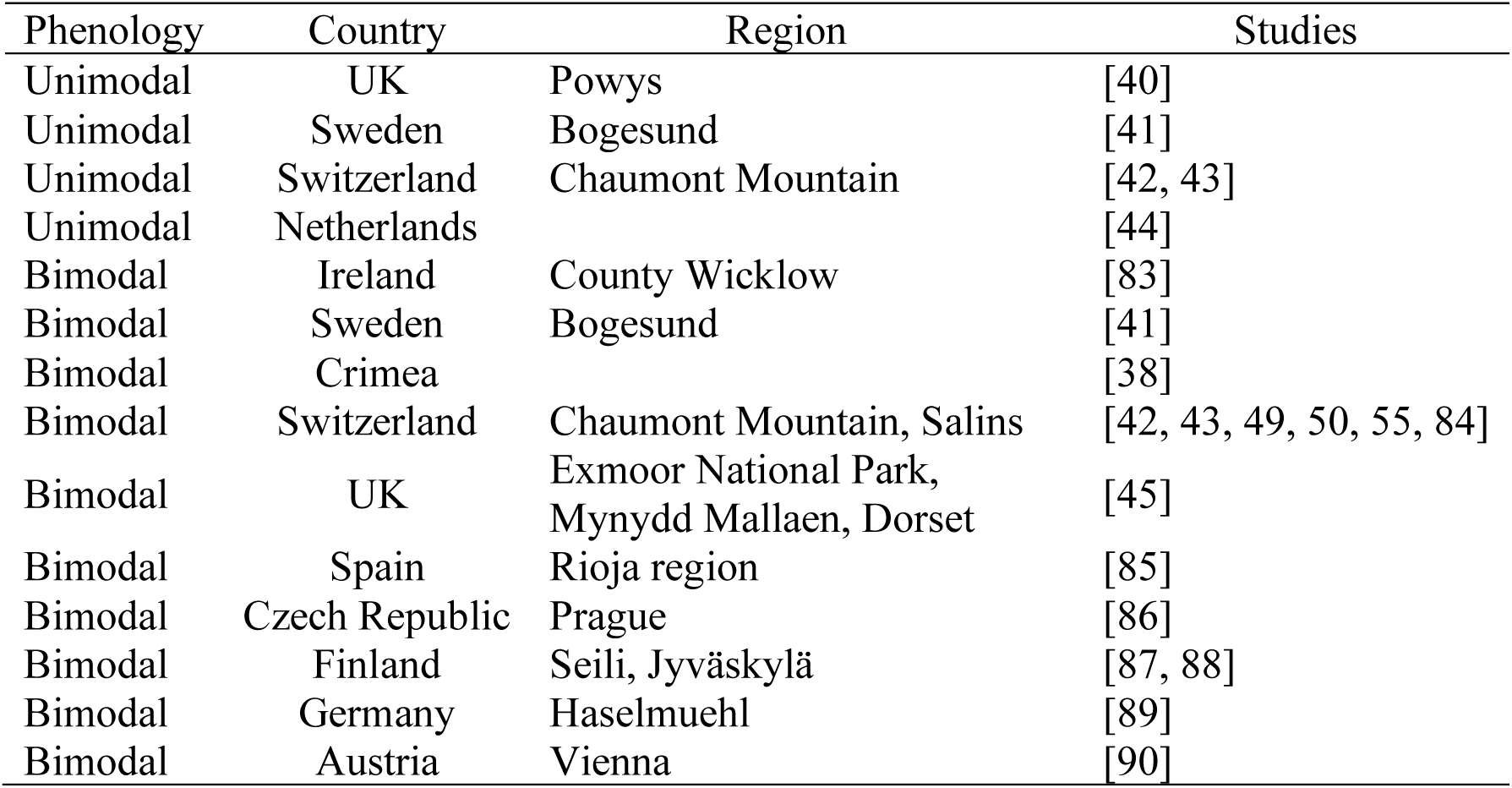
Type of phenology for Ixodes ricinus ticks in different countries in Europe. *Notes*: *Ixodes ricinus* ticks (nymphs and adults) either have a unimodal or bimodal phenology. The phenology is shown for different countries and regions in Europe. The decision of whether a study found a unimodal or bimodal phenology was based on whether the authors of the study identified the mode of phenology as either unimodal or bimodal in the article.

| Phenology | Country | Region | Studies |
| --- | --- | --- | --- |
| Unimodal | UK | Powys | [40] |
| Unimodal | Sweden | Bogesund | [41] |
| Unimodal | Switzerland | Chaumont Mountain | [42, 43] |
| Unimodal | Netherlands |  | [44] |
| Bimodal | Ireland | County Wicklow | [83] |
| Bimodal | Sweden | Bogesund | [41] |
| Bimodal | Crimea |  | [38] |
| Bimodal | Switzerland | Chaumont Mountain, Salins | [42, 43, 49, 50, 55, 84] |
| Bimodal | UK | Exmoor National Park,<br>Mynydd Mallaen, Dorset | [45] |
| Bimodal | Spain | Rioja region | [85] |
| Bimodal | Czech Republic | Prague | [86] |
| Bimodal | Finland | Seili, Jyväskylä | [87, 88] |
| Bimodal | Germany | Haselmuehl | [89] |
| Bimodal | Austria | Vienna | [90] |

The two alternative explanations for this bimodal phenology of questing *I. ricinus* nymphs are the developmental diapause hypothesis and the direct development hypothesis (Figure 1) [19, 38, 42, 43, 45–47]. The developmental diapause hypothesis (Figure 1) suggests that the timing of the larval blood meal splits the larval cohort into two groups of nymphs that are active in the spring and fall of the following year [18, 19]. Larvae that obtain their blood meal in early summer, moult into unfed nymphs, enter behavioral diapause in fall, overwinter as unfed nymphs, and quest the following spring. In contrast, larvae that obtain their blood meal in late summer, enter developmental diapause, overwinter as engorged larvae, complete their development the following summer, and quest the following fall [19]. The direct development hypothesis (Figure 1) suggests that the timing of the larval blood meal splits the larval cohort into two groups of nymphs that are active in the fall of that year and the spring of the following year [18, 19]. Larvae that obtain their blood meal in early summer, moult into unfed nymphs, and quest that same fall. In contrast, larvae that obtain their blood meal in late summer, moult into unfed nymphs, enter behavioral diapause, and quest the following spring [19]. In both hypotheses, there is a 1-year time lag between larval feeding and the spring nymphal peak. The critical distinction between these two hypotheses is the time lag between larval feeding and the fall nymphal peak, which is 1 year for the developmental diapause hypothesis and 0 years for the direct development hypothesis. To date, it is not clear which of these two hypotheses is more important for explaining the fall peak of *I. ricinus* nymphs.

**Figure 1.**
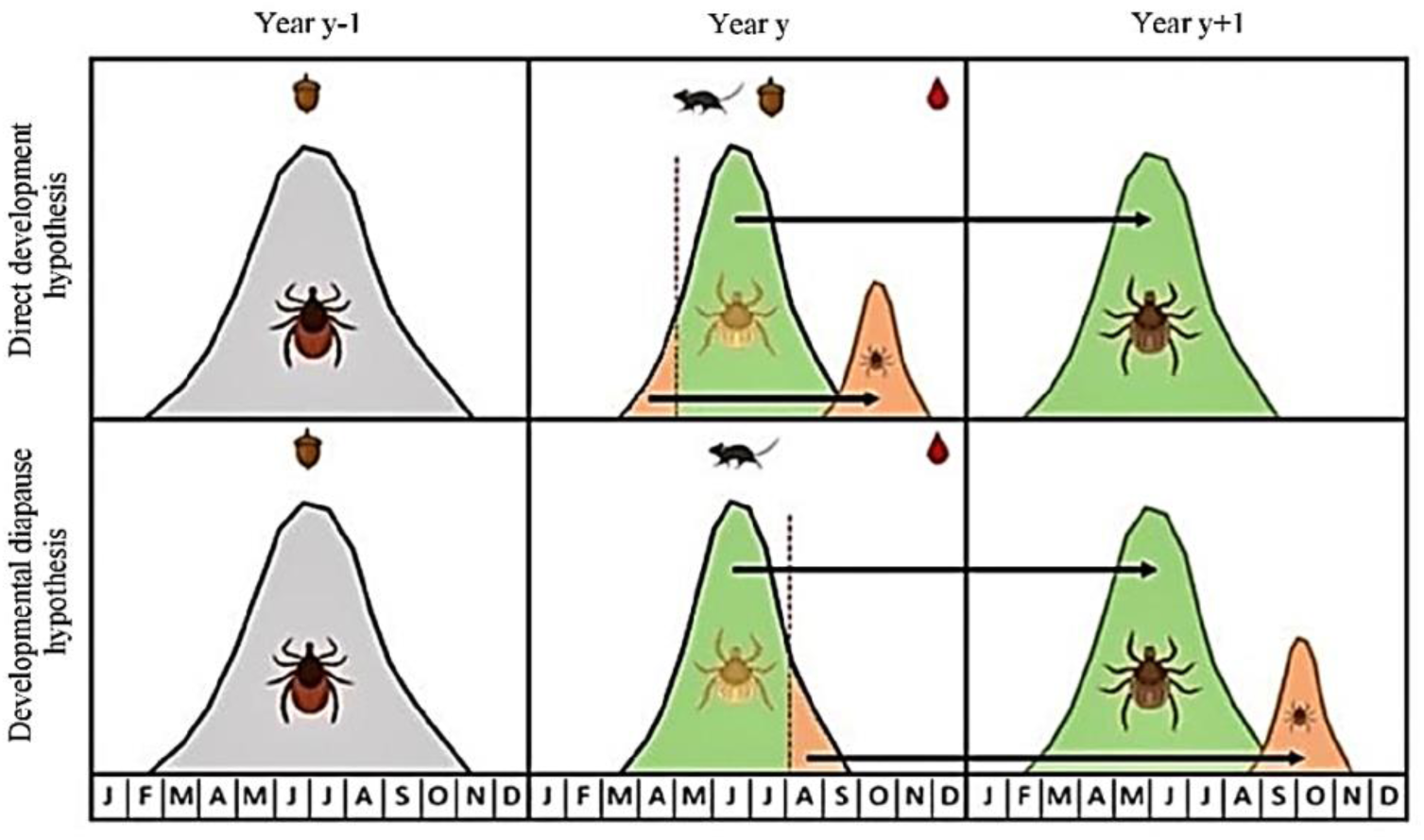
Two alternative (but not mutually exclusive) hypotheses for the fall peak are shown: the direct development hypothesis (top panel) and the developmental diapause hypothesis (bottom panel). The approximate 3-year life cycle of *I. ricinus* is shown by adult ticks (Year *y* – 1 in the left panel) that lay eggs, the larvae (Year *y* in the middle panel), and the nymphs (Year *y* + 1 in the right panel). The origin of the spring peak of nymphs (green peak in Year *y* + 1) is the same for both hypotheses and is as follows. Larvae that obtain their blood meal in the summer (green fraction in Year *y*), moult into unfed nymphs in the same year, enter behavioral diapause, overwinter, and quest as unfed nymphs the following spring (green peak in Year *y* + 1). The two hypotheses differ with respect to the origin of the fall nymphal peak (small orange peak). In the direct development hypothesis, larvae that obtain their blood meal early in the summer (orange fraction in Year *y*) moult into unfed nymphs in summer and quest as unfed nymphs that same fall (small orange peak in Year *y*). In the developmental diapause hypothesis, larvae that obtain their blood meal late in the summer (orange fraction in Year *y*), overwinter as engorged larvae, moult into unfed nymphs the following summer and quest as unfed nymphs the following fall (small orange peak in Year *y* + 1). The time lag between a masting event (Year *y* – 1) and the spring peak of nymphs (green peak in Year *y* + 1) is 2 years. In contrast, the time lag between a masting event and the fall peak of nymphs (small orange peak) is 1 year under the direct development hypothesis and 2 years under the developmental diapause hypothesis.

In addition to influencing the vital rates (development, survival, and reproduction) and the seasonal phenology, the climate also influences tick questing behavior, which in turn determines the probability that ticks are captured by common tick sampling methods (e.g., dragging). Thus, while the seasonal phenology in central Europe dictates that questing nymphs are most abundant in spring or early summer, the questing activity of nymphs on any given day in spring depends on the weather [21, 48, 49]. Field plot experiments have shown that the percentage of ticks that are actively questing depends on the weather [39, 50]. Questing activity is generally determined by the water balance regulation, which is affected by both temperature and relative humidity [51–53]. In summary, variation in the abundance of questing ticks at any given time depends on three separate mechanisms: (1) time-lagged ecological factors that influence the vital rates, (2) photoperiod-dependent diapause that determines the broad seasonal activity patterns of questing ticks, and (3) daily weather conditions interacting with the tick water balance that determine whether ticks will quest or not on that day. Separating these different mechanisms, which operate on different temporal scales, is not an easy task.

We have previously used a long-term data set (15 years) on the questing abundance of *I. ricinus* ticks at four different elevations on a mountain in western Switzerland to investigate the ecological factors that influence the inter-annual variation in the density of questing nymphs (DON) and the density of questing nymphs infected with the causative agents of Lyme borreliosis (DIN) [12, 36]. The most important finding in these two studies was that inter-annual variation in the DON and the DIN was strongly associated with inter-annual variation in the production of seeds by European beech trees 2 years prior. For these two studies, we analyzed the annual abundance of nymphs for the calendar year (January 1 to December 31). This approach assumes that the spring and fall peaks of questing nymphs in the same calendar year are both governed by the same ecological factors and time lags. This assumption is correct for the developmental diapause hypothesis, but incorrect for the direct development hypothesis. Another limitation of our previous study was that we investigated a highly limited set of climate variables calculated as annual means for either the current year or the previous year. In contrast, numerous studies suggest that season rather than calendar year is the relevant time scale over which climate variables influence the population ecology of *Ixodes* ticks [13, 54]. By dividing the calendar year into different seasons, we are increasing the temporal resolution at which our climate variables can explain inter-annual variation in tick abundance.

In the present study, we build on our previous modelling efforts of the same data set to investigate three objectives. First, determine whether the developmental diapause hypothesis or the direct development hypothesis is better at explaining inter-annual variation in the fall peak of nymphs. Second, determine whether seasonal climate means are better than annual climate means at predicting inter-annual variation in the density of nymphs, and which seasonal climate variables are important. Third, determine whether we can use generalized additive models (GAMs) to model the complex bimodal seasonal phenology of *I. ricinus* nymphs and whether this approach yields additional insights into the factors that explain seasonal variation in questing tick abundance.

## METHODS

### Study location

The study was conducted on the south-facing slope of Chaumont Mountain, which is part of the Jura mountains, and is in the canton of Neuchâtel, in western Switzerland. Four tick sampling sites, referred to as low, medium, high, and top, were established at elevations of 620, 740, 900, and 1073 m above sea level (ASL), respectively, and have been described previously [43, 55, 56]. There is logging in the area, and there are hiking trails and recreation areas for the public. The forest on Chaumont Mountain is mainly composed of European beech (*Fagus sylvatica*; 28.6%), Norway spruce (*Picea abies*; 28.5%), European silver fir (*Abies alba*; 20.4%), sycamore maple (*Acer pseudoplatanus*; 5.9%), European ash (*Fraxinus excelsior*; 3.7%), Scots pine (*Pinus sylvestris*; 2.3%), sessile oak (*Quercus petraea*; 2.3%), willow (*Salix* spp.; 2.1%), common whitebeam (*Sorbus aria*; 1.6%), and European hornbeam (*Carpinus betulus*; 1.0%) [57].

### Sampling *I. ricinus* ticks in the field

Questing *I. ricinus* nymphs and adult ticks were collected monthly over a period of 15 years (January 2004 to December 2018) at each of the four elevation sites. The sampling protocol has been described previously [55]. Briefly, a 1-m^2^ cotton flag was dragged over the ground along a fixed transect with a length of 100 m (low elevation) or 120 m (other elevations). The flag was inspected every 20 m and nymphs and adult ticks were counted separately and placed in collection vials. This method of tick collection targets questing ticks and removes them from the environment; these removed ticks cannot be encountered on future sampling occasions and they cannot contribute to future tick population growth. The same person (Olivier Rais) conducted all 720 transects (4 elevations*15 years*12 months = 720 transects). No dragging was performed on days when there was snow on the ground (hereafter referred to as snow days). Over the study period, a total of 34 snow days occurred, which resulted in missing data for 136 transects.

### Field-collected climate variables

Temperature (units are °C) and relative humidity (RH; units are %) were recorded at 60 cm above ground at one moment in time on the day of tick collection (usually between 10:00 am and 2:00 pm) at each tick sampling site using a thermohygrometer (Model 615, Testo SA, Lonay, Switzerland). Thus, for each combination of elevation and year, we had a total of 12 field-collected measurements of temperature and RH. The saturation deficit (SD) is a measure of the drying power of the atmosphere (units are mm of mercury) and is calculated using temperature (T; units are °C) and RH (units are %) as follows: SD = (1 − RH/100) * 4.9463 * e^0.0621T^ [42, 58]. The accuracy of our field-collected climate data was confirmed by comparing them to the weather station data [12].

### Weather station climate variables

We also obtained climate data from the CLIMAP-net database of the Federal Office of Meteorology and Climatology MeteoSwiss. Two weather stations close to our study site are in Neuchâtel at 485 m ASL (WMO number = 06604) and in Chaumont at 1136 m ASL (WMO number = 06608). These weather stations sample the temperature and RH every hour, and the total precipitation each day at 200 cm above ground. We used the daily mean temperature (average of the 24 measurements per day), the daily mean RH (average of the 24 measurements per day), and the daily total precipitation (rain, snow). Thus, for each year, we had a total of 365 weather station measurements of these three climate variables. The SD was calculated as previously described. For each of the four elevation sites, we calculated site-specific climate variables by interpolating the data between the two weather stations (Additional file 1: Section 1).

### Data on inter-annual variation in tree masting

We previously demonstrated that the abundance of *I. ricinus* ticks depends on the seed production of deciduous trees [12, 36]. The seeds or fruit of forest trees (e.g., acorns of oak trees or the beech nuts of beech trees) are often referred to as mast. The annual production of mast by a population of trees in an area occurs in the fall and is highly variable among years [59]. The MASTREE database contains data on masting (or seed production) for many locations in Europe from year 1982 to year 2016 for two tree species, European beech (*Fagus sylvatica*) and Norway spruce (*Picea abies*) [60]. In this database, the mast intensity is classified into five classes: 1, 2, 3, 4, and 5, which refer to very poor mast, poor mast, moderate mast, good mast, and full mast, respectively [60]. We used the MASTREE database [60] to obtain masting data for the European beech and Norway spruce for the canton of Neuchâtel for the years of our study. These two species of tree account for 57.1% of the trees at our study location.

### Fat content of *I. ricinus* nymphs collected in the spring and fall

Fat is a non-renewable source of energy derived from each blood meal that ticks use to quest for hosts and to maintain their water balance [58, 61, 62]. As *I. ricinus* feeds once per life stage and has no other energy sources between blood meals, their fat content is an index of their current age and future longevity in the unfed state [40, 45, 58]. In a previous study on the *I. ricinus* population at our field site, we had collected nymphs in the spring and fall of 2010 and measured their fat content [63]. In the present study, we compared the fat content between the spring and fall nymphs. The direct development hypothesis predicts that the fall nymphs are younger and should therefore have a higher fat content compared to the spring nymphs (∼3 months versus ∼9 months since the larval blood meal). In contrast, the delayed diapause hypothesis predicts that the fall nymphs are older and should therefore have a lower fat content compared to the spring nymphs.

### Statistical methods

The MASTREE database contains data on masting from year 1982 to year 2016, whereas our tick surveillance study ran from January 2004 to December 2018. We therefore had the beech masting scores one year prior for the DON in 2017 but not for the DON in 2018. For this reason, the statistical analyses are restricted to a 14-year period (2004 to 2017).

#### The density of nymphs (DON)

The density of nymphs (DON) is a measure of the monthly abundance of questing nymphs per 100 m^2^. The density of infected nymphs (DIN) is the number of questing nymphs infected with *B. burgdorferi* sl per 100 m^2^. Over the 14-year study period that was covered by the MASTREE database, estimates of the DON and DIN were obtained from 558 fixed monthly transects (4 elevations * 14 years * 12 months = 672 transects; 114 missing transects due to snow and other reasons; 672 transects – 114 transects = 558 transects). The transects on the snow days were coded as missing data.

#### Definition of the spring nymphal peak and the fall nymphal peak

Previous work on the abundance of questing *I. ricinus* nymphs at Chaumont Mountain and at other nearby sites have shown a bimodal phenology with a large peak of the DON in the spring and a smaller peak of the DON in the fall [42, 43, 49, 50, 55]. To test which variables are best at explaining inter-annual variation in the spring and fall peaks of the DON, a date must be chosen to separate these two groups of nymphs. We decided that the spring peak included the nymphs sampled from January 1 to August 31, whereas the fall peak included the nymphs sampled from September 1 to December 31. This date was chosen because the DON reached a minimum at this time.

#### Weather on the day of tick sampling

To investigate if the weather influenced nymphal questing activity, we used the field-measured weather data on the day of tick sampling. An important advantage of the field-measured data compared to the weather station data was that they are specific for each of the four elevation sites. We did not use the field-measured weather data to calculate annual or seasonal means because there were not enough data (i.e., only 12 measurements per calendar year at each site).

#### Annual and seasonal mean climate variables

Life history traits (development, survival, reproduction) of tick populations depend on abiotic factors such as temperature, RH, SD, and precipitation (rain, snow). A great unknown is the relevant time frame over which these abiotic climate variables affect the vital rates of tick populations. For example, the DON in the spring might depend on the climate conditions of the previous winter (e.g., overwinter survival of nymphs), or on the climate conditions of the previous summer, which would influence the rates at which larvae obtain and digest their blood meals, and moult into nymphs. In our previous studies on inter-annual variation in the DON and DIN [12, 36], we calculated annual means for the climate variables that were based on a 12-month calendar year. However, many steps in the tick life cycle happen over shorter time scales; for example, engorged larvae take ∼ 6 to 8 weeks to moult into nymphs at room temperature [64]. Thus, the life history traits of tick populations may depend on climate variables that are operating over shorter temporal windows (e.g., seasons rather than years). We therefore calculated mean seasonal climate variables for each of the 12 seasons (3 years * 4 seasons per year = 12 seasons) that preceded and encompassed the year of tick collection (Figure 2). The seasonal means for the winter, spring, summer, and fall were calculated as follows: December 1 (e.g., previous year) to February 28/29, March 1 to May 31, June 1 to August 31, and September 1 to November 30. As time lags are important in tick ecology, we calculated our annual mean climate variables and seasonal mean climate variables in the present year, the previous year, or two years prior (Figure 2). Thus, for each climate variable, there were a total of 12 seasonal means (3 years * 4 seasons per year = 12) and 3 annual means. The exception was the annual snow fall (units of cm) which was calculated over the time from October 1 (e.g., previous year) to May 31.

**Figure 2.**
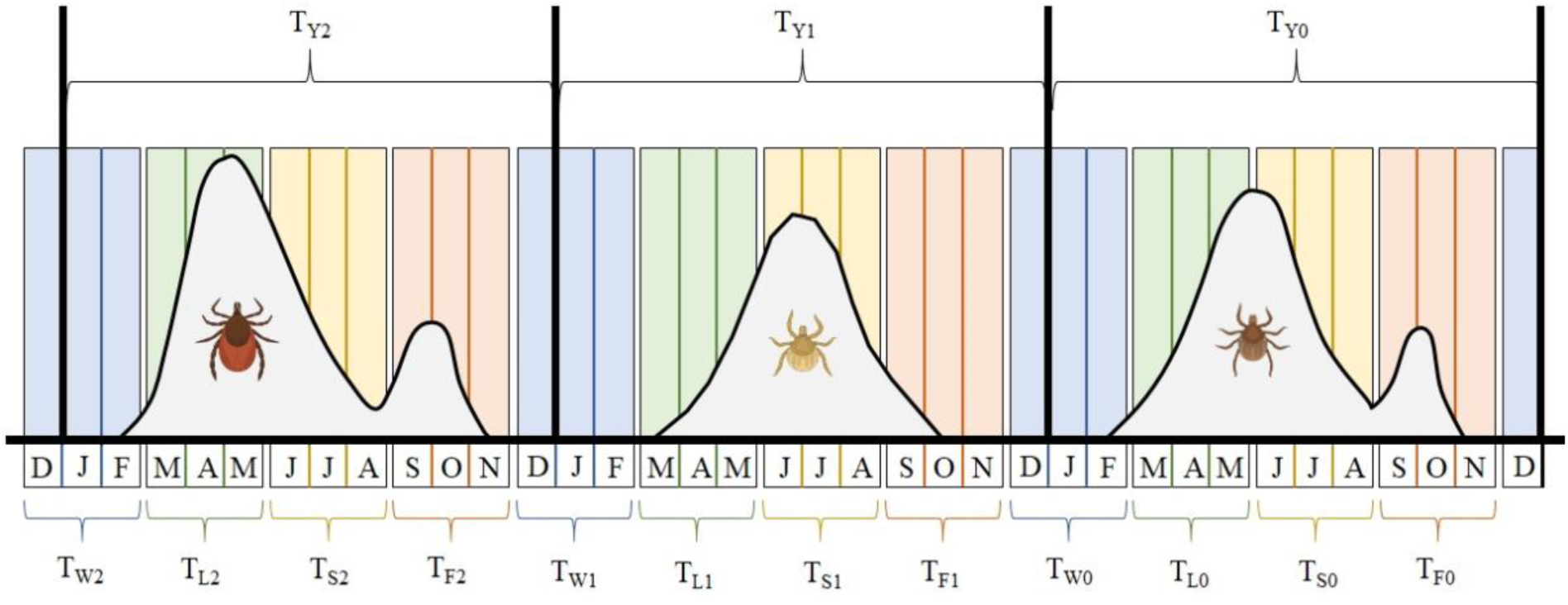
Visual representation of how the mean annual climate variables and the mean seasonal climate variables were calculated over 3 different years. The approximate 3-year life cycle of *I. ricinus* is shown by adult ticks that lay eggs (left panel), the eggs that hatch into larvae (middle panel), and the larvae that become nymphs (right panel). The bimodal or unimodal distribution in each panel represents the phenology of each tick stage. The three calendar years are shown by the vertical black lines, the months are shown by the bars that are labelled below the X-axis. The four seasons of winter, spring, summer, and fall are color-coded as blue, green, yellow, and orange, respectively. The response variable of interest is the density of nymphs (DON) in the right panel. For illustrative purposes, the explanatory variable is temperature (abbreviated as T). The 3 annual means of the temperature are T_Y2_, T_Y1_, and T_Y0_ (labelled at the top); the subscript ‘Y’ indicates that the temperature is averaged over the calendar year; the subscripts 0, 1, and 2 indicate the time lag (i.e., 0, 1, or 2 years before the year of the DON). There are 12 seasonal means of the temperature (labelled at the bottom); the subscripts ‘W’, ‘L’, ‘S’, and ‘F’ indicate that the temperature is averaged over the winter, lent (spring), summer, and fall; the subscripts 0, 1, and 2 indicate the time lag in years. For example, T_W2_ is the mean temperature of the winter two years before the year of the DON.

#### Annual beech masting score

Previous studies have shown that there is a 2-year time lag between masting events and the annual DON and the annual DIN [12, 14, 35, 36]. Our recent analyses of the same data showed that inter-annual variation in the DON and the DIN was strongly associated with the mast scores 2 years prior of European beech but not Norway spruce [12, 36]. Upon further reflection, we realized that while the 2-year time lag is true for the spring peak it may not be true for the fall peak. The developmental diapause hypothesis and the direct development hypothesis predict that the time lag between the beech masting score and the fall peak of nymphs should be 2 years versus 1 year, respectively. To test these two hypotheses, we created three different explanatory variables, BM_2/2_, BM_1/1_, and BM_2/1_, for the beech masting (BM) score. BM_2/2_ assumes there is a 2-year time lag between BM and the spring and fall peaks of nymphs. BM_1/1_ assumes there is a 1-year time lag between BM and the spring and fall peaks of nymphs. BM_2/1_ assumes there is a 2-year time lag and a 1-year time lag between BM and the spring and fall peak, respectively. BM_2/2_ is consistent with the developmental diapause hypothesis, BM_2/1_ is consistent with the direct development hypothesis, and BM_1/1_ is not consistent with either hypothesis.

#### The monthly DON was analyzed using generalized additive models (GAMs)

We used generalized additive models (GAMs) with extra-binomial errors to model the non-linear bimodal seasonal phenology of the monthly DON, which represent count data. GAMs are like generalized linear models, but an important difference is that smoother functions are used to model the response variable as a non-linear function of one or more explanatory variables. The smoother functions are non-parametric (they are like moving average functions) and can be used to fit any complex curve and this flexibility is a great strength of the GAM approach. A weakness is that there are no parameter estimates for the explanatory variables that are modelled with the smoother functions, which makes model interpretation more difficult.

#### Parametric versus non-parametric functions of explanatory variables

The major motivation for using GAMS was to capture the bimodal non-linear seasonal phenology of the DON over the calendar year. We compared the ability of smoother functions of calendar day, temperature, RH, and SD to capture this seasonal phenology. We also used the smoother function as a tool to investigate which explanatory variables were important for explaining variation in the DON without having to specify the nature of the relationship. However, if the relationship between the DON and an explanatory variable was obviously linear or quadratic, then we replaced the non-parametric smoother function of this explanatory variable with a parametric function because model interpretation is easier for simple parametric functions compared to complex non-parametric functions.

#### Variables used to model the DON

The explanatory variables that were investigated included elevation site (4 levels: low, medium, high, top), the covariate year (rescaled as 1, 2, 3, …, 14), the covariate day (rescaled as 1, 2, 3, …, 365), the covariate beech mast score (range: 1 to 5), the field-collected climate variables of temperature (t), relative humidity (rh), and saturation deficit (sd) on the day of tick sampling, and the mean annual and mean seasonal climate variables obtained from the weather stations for temperature (T), RH, SD, precipitation (PR), and annual snow fall (SN). For beech mast score there were three different types of variables, BM_2,2_, BM_1,1_, and BM_2,1_, as previously discussed. For 4 climate variables (T, RH, SD, PR), there were 3 annual means and 12 seasonal means; for annual SN there were 3 annual means. The 72 explanatory variables and their acronyms are listed in Table 2.

**Table 2.** Acronym and definition of each variable used in the present study.

| Acronym | Description |
| --- | --- |
| DON | Monthly density of nymphs per 100m<sup>2</sup> |
| S | Site name (factor with 4 levels: low, medium, high, top) |
| Y | Year of the study (covariate: 1, 2, ..., 15) |
| D | Day of the year for the tick sampling (covariate: 1, 2, ..., 365) |
| BM<sub>1/1</sub> | Beech mast score in year y-1 (covariate: 1, 2, ..., 5) |
| BM<sub>2/2</sub> | Beech mast score in year y-2 (covariate: 1, 2, ..., 5) |
| BM<sub>2/1</sub> | Beech mast score with a 2-year time lag for the spring peak and a 1-year time lag for the fall peak (covariate: 1, 2, ..., 5) |
| t | Temperature on day of sampling from the field-collected data (°C) |
| rh | Relative humidity on day of sampling from the field-collected data (%) |
| sd | Saturation deficit on day of sampling from the field-collected data (mmHg) |
| T<sub>Y0</sub> | Mean annual temperature in year y from the weather station data (°C) |
| T<sub>Y1</sub> | Mean annual temperature in year y-1 from the weather station data (°C) |
| T<sub>Y2</sub> | Mean annual temperature in year y-2 from the weather station data (°C) |
| RH<sub>Y0</sub> | Mean annual relative humidity in year y from the weather station data (%) |
| RH<sub>Y1</sub> | Mean annual relative humidity in year y-1 from the weather station data (%) |
| RH<sub>Y2</sub> | Mean annual relative humidity in year y-2 from the weather station data (%) |
| SD<sub>Y0</sub> | Mean annual saturation deficit in year y from the weather station data (mmHg) |
| SD<sub>Y1</sub> | Mean annual saturation deficit in year y-1 from the weather station data (mmHg) |
| SD<sub>Y2</sub> | Mean annual saturation deficit in year y-2 from the weather station data (mmHg) |
| PR<sub>Y0</sub> | Mean annual precipitation in year y from the weather station data (mm) |
| PR<sub>Y1</sub> | Mean annual precipitation in year y-1 from the weather station data (mm) |
| PR<sub>Y2</sub> | Mean annual precipitation in year y-2 from the weather station data (mm) |
| SN<sub>Y0</sub> | Annual snowfall in year y from the weather station data (cm) |
| SN<sub>Y1</sub> | Annual snowfall in year y-1 from the weather station data (cm) |
| SN<sub>Y2</sub> | Annual snowfall in year y-2 from the weather station data (cm) |
| T<sub>F0</sub> | Mean seasonal temperature of the fall in year y from the weather station data (°C; mean daily temperature was averaged over months of September, October, and November) |
| T<sub>S0</sub> | Mean seasonal temperature of the summer in year y from the weather station data (°C; mean daily temperature was averaged over months of June, July, and August) |
| T<sub>L0</sub> | Mean seasonal temperature of the spring in year y from the weather station data (°C; mean daily temperature was averaged over months of March, April, and May) |
| T<sub>W0</sub> | Mean seasonal temperature of the winter in year y from the weather station data (°C; mean daily temperature was averaged over months of December, January, and February) |
| T<sub>F1</sub> | Mean seasonal temperature of the fall in year y-1 from the weather station data (°C; mean daily temperature was averaged over months of September, October, and November) |
| T<sub>S1</sub> | Mean seasonal temperature of the summer in year y-1 from the weather station data (°C; mean daily temperature was averaged over months of June, July, and August) |
| T<sub>L1</sub> | Mean seasonal temperature of the spring in year y-1 from the weather station data (°C; mean daily temperature was averaged over months of March, April, and May) |
| T<sub>W1</sub> | Mean seasonal temperature of the winter in year y-1 from the weather station data (°C; mean daily temperature was averaged over months of December, January, and February) |
| T<sub>F2</sub> | Mean seasonal temperature of the fall in year y-2 from the weather station data (°C; mean daily temperature was averaged over months of September, October, and November) |
| T<sub>S2</sub> | Mean seasonal temperature of the summer in year y-2 from the weather station data (°C; mean daily temperature was averaged over months of June, July, and August) |
| T<sub>L2</sub> | Mean seasonal temperature of the spring in year y-2 from the weather station data (°C; mean daily temperature was averaged over months of March, April, and May) |
| T<sub>W2</sub> | Mean seasonal temperature of the winter in year y-2 from the weather station data (°C; mean daily temperature was averaged over months of December, January, and February) |
| RH<sub>F0</sub> | Mean seasonal relative humidity of the fall in year y from the weather station data (%; mean daily relative humidity was averaged over months of September, October, and November) |
| RH<sub>S0</sub> | Mean seasonal relative humidity of the summer in year y from the weather station data (%; mean daily relative humidity was averaged over months of June, July, and August) |
| RH<sub>L0</sub> | Mean seasonal relative humidity of the spring in year y from the weather station data (%; mean daily relative humidity was averaged over months of March, April, and May) |
| RH<sub>W0</sub> | Mean seasonal relative humidity of the winter in year y from the weather station data (%; mean daily relative humidity was averaged over months of December, January, and February) |
| RH<sub>F1</sub> | Mean seasonal relative humidity of the fall in year y-1 from the weather station data (%; mean daily relative humidity was averaged over months of September, October, and November) |
| RH<sub>S1</sub> | Mean seasonal relative humidity of the summer in year y-1 from the weather station data (%; mean daily relative humidity was averaged over months of June, July, and August) |
| RH<sub>L1</sub> | Mean seasonal relative humidity of the spring in year y-1 from the weather station data (%; mean daily relative humidity was averaged over months of March, April, and May) |
| RH<sub>W1</sub> | Mean seasonal relative humidity of the winter in year y-1 from the weather station data (%; mean daily relative humidity was averaged over months of December, January, and February) |
| RH<sub>F2</sub> | Mean seasonal relative humidity of the fall in year y-2 from the weather station data (%; mean daily relative humidity was averaged over months of September, October, and November) |
| RH<sub>S2</sub> | Mean seasonal relative humidity of the summer in year y-2 from the weather station data (%; mean daily relative humidity was averaged over months of June, July, and August) |
| RH<sub>L2</sub> | Mean seasonal relative humidity of the spring in year y-2 from the weather station data (%; mean daily relative humidity was averaged over months of March, April, and May) |
| RH<sub>W2</sub> | Mean seasonal relative humidity of the winter in year y-2 from the weather station data (%; mean daily relative humidity was averaged over months of December, January, and February) |
| SD<sub>F0</sub> | Mean seasonal saturation deficit of the fall in year y from the weather station data (mmHg; mean daily saturation deficit was averaged over months of September, October, and November) |
| SD<sub>S0</sub> | Mean seasonal saturation deficit of the summer in year y from the weather station data (mmHg; mean daily saturation deficit was averaged over months of June, July, and August) |
| SD<sub>L0</sub> | Mean seasonal saturation deficit of the spring in year y from the weather station data (mmHg; mean daily saturation deficit was averaged over months of March, April, and May) |
| SD<sub>W0</sub> | Mean seasonal saturation deficit of the winter in year y from the weather station data (mmHg; mean daily saturation deficit was averaged over months of December, January, and February) |
| SD<sub>F1</sub> | Mean seasonal saturation deficit of the fall in year y-1 from the weather station data (mmHg; mean daily saturation deficit was averaged over months of September, October, and November) |
| SD<sub>S1</sub> | Mean seasonal saturation deficit of the summer in year y-1 from the weather station data (mmHg; mean daily saturation deficit was averaged over months of June, July, and August) |
| SD<sub>L1</sub> | Mean seasonal saturation deficit of the spring in year y-1 from the weather station data (mmHg; mean daily saturation deficit was averaged over months of March, April, and May) |
| SD<sub>W1</sub> | Mean seasonal saturation deficit of the winter in year y-1 from the weather station data (mmHg; mean daily saturation deficit was averaged over months of December, January, and February) |
| SD<sub>F2</sub> | Mean seasonal saturation deficit of the fall in year y-2 from the weather station data (mmHg; mean daily saturation deficit was averaged over months of September, October, and November) |
| SD<sub>S2</sub> | Mean seasonal saturation deficit of the summer in year y-2 from the weather station data (mmHg; mean daily saturation deficit was averaged over months of June, July, and August) |
| SD<sub>L2</sub> | Mean seasonal saturation deficit of the spring in year y-2 from the weather station data (mmHg; mean daily saturation deficit was averaged over months of March, April, and May) |
| SD<sub>W2</sub> | Mean seasonal saturation deficit of the winter in year y-2 from the weather station data (mmHg; mean daily saturation deficit was averaged over months of December, January, and February) |
| PR<sub>F0</sub> | Mean seasonal precipitation of the fall in year y from the weather station data (mm; mean daily precipitation was averaged over months of September, October, and November) |
| PR<sub>S0</sub> | Mean seasonal precipitation of the summer in year y from the weather station data (mm; mean daily precipitation was averaged over months of June, July, and August) |
| PR<sub>L0</sub> | Mean seasonal precipitation of the spring in year y from the weather station data (mm; mean daily precipitation was averaged over months of March, April, and May) |
| PR<sub>W0</sub> | Mean seasonal precipitation of the winter in year y from the weather station data (mm; mean daily precipitation was averaged over months of December, January, and February) |
| PR<sub>F1</sub> | Mean seasonal precipitation of the fall in year y-1 from the weather station data (mm; mean daily precipitation was averaged over months of September, October, and November) |
| PR<sub>S1</sub> | Mean seasonal precipitation of the summer in year y-1 from the weather station data (mm; mean daily precipitation was averaged over months of June, July, and August) |
| PR<sub>L1</sub> | Mean seasonal precipitation of the spring in year y-1 from the weather station data (mm; mean daily precipitation was averaged over months of March, April, and May) |
| PR<sub>W1</sub> | Mean seasonal precipitation of the winter in year y-1 from the weather station data (mm; mean daily precipitation was averaged over months of December, January, and February) |
| PR<sub>F2</sub> | Mean seasonal precipitation of the fall in year y-2 from the weather station data (mm; mean daily precipitation was averaged over months of September, October, and November) |
| PR<sub>S2</sub> | Mean seasonal precipitation of the summer in year y-2 from the weather station data (mm; mean daily precipitation was averaged over months of June, July, and August) |
| PR<sub>L2</sub> | Mean seasonal precipitation of the spring in year y-2 from the weather station data (mm; mean daily precipitation was averaged over months of March, April, and May) |
| PR<sub>W2</sub> | Mean seasonal precipitation of the winter in year y-2 from the weather station data (mm; mean daily precipitation was averaged over months of December, January, and February) |

#### Count data and extra-binomial errors

As the DON represents count data (number of nymphs per 100 m^2^), we initially used the Poisson distribution to model the residual errors. The residuals from these models were highly overdispersed; the ratio of the residual deviance (7025) to the residual degrees of freedom (500) was > 1 (7025/500 = 14.05). This problem of overdispersion was solved by using the negative binomial distribution to model the residual errors; the residuals from these models were no longer overdispersed (517/514 = 1.01). Parameter estimates from models with Poisson errors (or extra-binomial errors) are reported on a log scale. Thus, the exponential function must be applied to the parameter estimates to determine the effect size of the explanatory variables on the DON on the original scale.

Models with Poisson errors (or extra-binomial errors) analyze count data, which are integers. However, the monthly DONs were not always integers because they had been measured over a transect area of 120 m^2^ before being standardized to an area of 100 m^2^. These non-integer DON values had to be rounded to the nearest integer to run GAMs with Poisson errors (or extra-binomial errors).

#### Scaling of the explanatory climate variables

All the annual and seasonal climate variables were transformed to z-scores with a mean of 0 and a standard deviation of 1. This transformation facilitates the comparison of slopes between climate variables measured in different units. One problem with modelling the response variable as a quadratic relationship of the explanatory variable is the strong collinearity between the linear and quadratic terms of the explanatory variable. To solve this problem, we transformed the explanatory variables using the *poly()* function in R so that the linear and quadratic terms were orthogonal to each other.

#### Model selection approach

To identify the best model, we used a model selection approach based on the Akaike information criterion (AIC). Models were ranked according to their AIC values, and the Akaike weights, which indicate the percent support, were calculated for each model. We used the Akaike weights to calculate the model-averaged parameter estimates and their 95% confidence intervals (CIs). For the generalized additive models that analyzed the DON, we assessed the goodness of fit for the best model from the model selection table (Additional file 1: Section 2).

#### Statistical software

We used R version 4.0.3 for all statistical analyses [65]. We used the *gam()* function in the mgcv package to run the GAMs [66]. We used the *poly()* function in the base package to rescale the linear and quadratic terms of each explanatory climate variable to avoid problems with collinearity. We used the *mod.sel()* function and the *model.av()* function in the MuMIn package to create the model selection tables and the model-averaged parameter estimates [67]. We used the *ggplot()* function in the ggplot2 package to create the graphs that show the effect sizes of the explanatory variables [68].

## RESULTS

### Mean monthly DON at each of the four elevation sites

The mean monthly DON was inversely related to the altitudinal gradient; it was highest at the low elevation site and lowest at the top elevation site (Table 3; Additional file 1: Section 3). The mean monthly DON at the low, medium, high, and top elevation sites was 74.4, 61.4, 42.7, and 10.6 nymphs per 100 m^2^, respectively (Table 3). If the low elevation site is set as the reference, the mean monthly DON at the medium, high, and top elevation sites were 19.3%, 41.2%, and 87.1% lower, respectively (Table 3; Additional file 1: Section 3). These estimates of the mean DON do not consider the effects of any other explanatory variables.

**Table 3.** Monthly density of nymphs (DON) per 100 m^2^ for the four elevation sites on Chaumont Mountain over the 15 years of the study (2004 – 2018). *Notes:* Shown for each of the four elevation sites are the number of years, sample size (N; units are number of transects), mean (units are nymphs per 100 m^2^), standard deviation, and the range of the monthly DON. For each elevation site, the expected sample size is 180 transects; missing transects are due to snow days when dragging for ticks was not possible. The DON is biased high due to the missing values for the snow days. In our previous study [12], we calculated the cumulative nymphal density (CND) for each calendar year by integrating the area under the curve of the seasonal phenology of the DON (per 100 m^2^) from January 1 to December 31. When this CND is divided by 365 days, it gives a second estimate of density of nymphs (DON2) that is less biased by the missing snow days. For this reason, the estimates of DON2 are lower than the DON.

| Site | Years | N | DON Mean | DON StdDev | DON Range | CND | DON2 |
| --- | --- | --- | --- | --- | --- | --- | --- |
| Low | 15 | 144 | 74.4 | 82.6 | 0.00 – 430.0 | 22,629 | 62.0 |
| Medium | 15 | 144 | 61.4 | 75.0 | 0.00 – 385.8 | 18,849 | 51.6 |
| High | 15 | 137 | 42.7 | 52.0 | 0.00 – 257.5 | 12,582 | 34.5 |
| Top | 15 | 133 | 10.6 | 14.5 | 0.00 – 80.8 | 3,000 | 8.2 |

### *I. ricinus* nymphs have a bimodal phenology on Chaumont Mountain

The seasonal changes in the DON over the calendar year at the four elevation sites clearly showed a bimodal phenology with a large peak of the DON in the spring and a smaller peak of the DON in the fall (Figure 3). This bimodal phenology of the nymphs was observed at the low, medium, and high elevation sites, but not at the top elevation site where the phenology was characterized by a single peak in the spring. For each of the four elevation sites, we compared the size of the spring and fall nymphal peaks by calculating the area under the curve of the seasonal phenology (Table 4). Expressed as a percent of the total, the fall peak at the low, medium, high, and top elevation sites was 14.9%, 11.5%, 12.5%, and 5.5%, respectively. Thus, the fall peak was largest for the low site and smallest for the top site. Interestingly, the spring peak occurred in April for the low site whereas it occurred in May at the medium, high, and top elevation sites.

**Figure 3.**
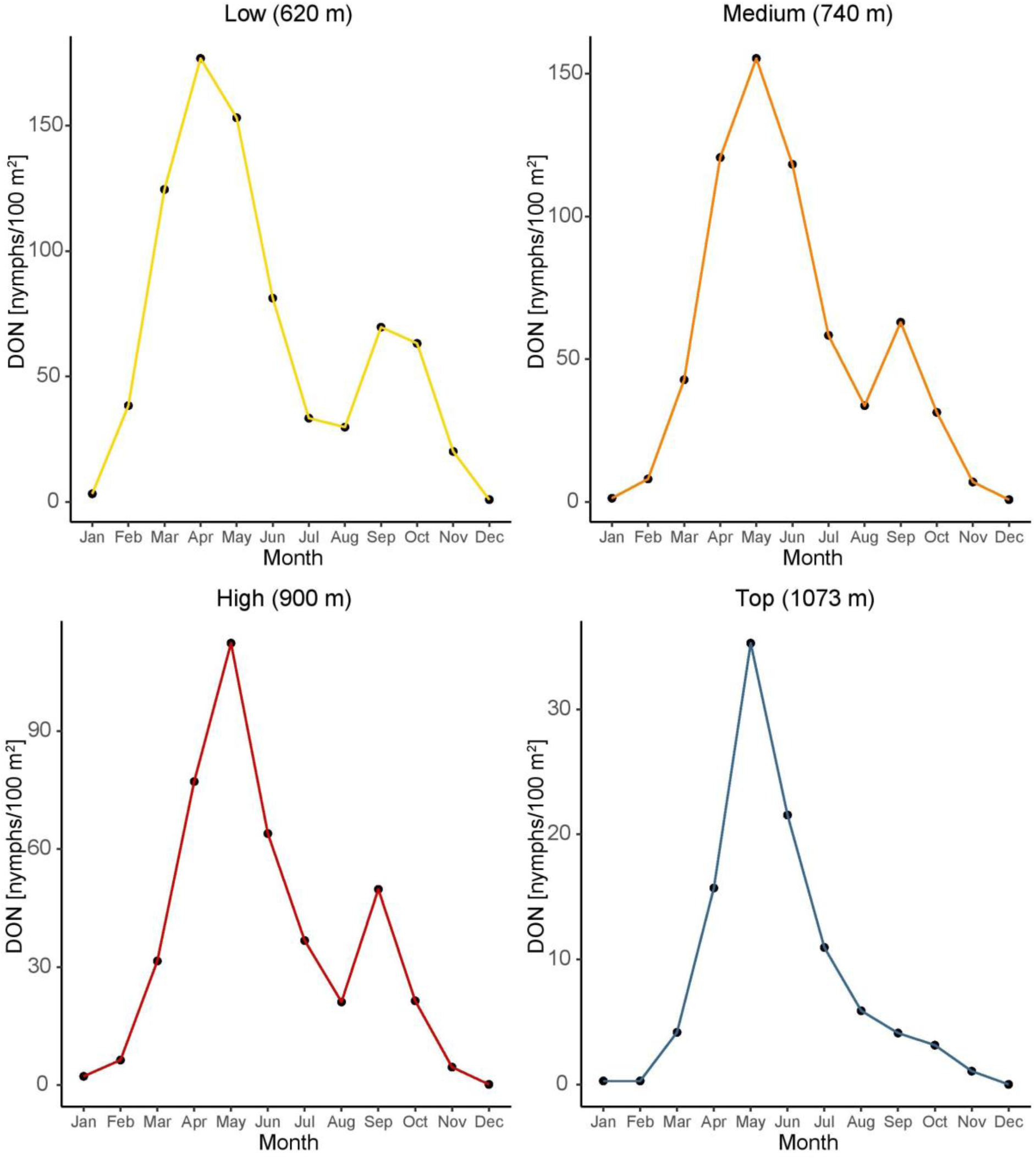
Seasonal changes in the DON over the calendar year at the four elevation sites. The DON is an estimate of the number of questing *I. ricinus* nymphs per 100 m^2^ sampled by the dragging method each month. For each month, the data are averaged over the 15 years of the study (2004 – 2018). A bimodal phenology with a large peak of the DON in the spring and a smaller peak of the DON in the fall was observed at the low, medium, and high elevation sites, but not at the top elevation site where the phenology was characterized by a single peak in the spring.

**Table 4.** Size of the spring and fall peaks of I. ricinus nymphs for the four elevation sites on Chaumont Mountain. Notes: The size of the spring peak and the fall peak of *I. ricinus* nymphs are shown for each of the four elevation sites on Chaumont Mountain. To compare the size of the cumulative spring peak (CSP) and the cumulative fall peak (CFP), we integrated the area under the curve of the seasonal phenology of the DON (per 100 m^2^) from January 1 to August 31 (CSP), and from September 1 to December 31 (CFP), respectively. The interpretation of the CSP and CFP are the numbers of *I. ricinus* nymphs that would have been captured if we had sampled for ticks every day over the corresponding calendar dates. For the CSP and the CFP, the sample size (N = 15 years), mean, standard deviation (StdDev), and range are shown. A second estimate of the cumulative nymphal density (CND2) was calculated by summing the CSP and the CFP. To express the two peaks as a percent, the CSP and the CFP were each divided by the CND2.

| Cumulative spring peak (CSP) |  |  |  |  |  |  |
| --- | --- | --- | --- | --- | --- | --- |
| Site | N | CSP Mean | CSP StdDev | CSP Range | CND2 | CSP (%) |
| Low | 15 | 18,145 | 10,508 | 7,516 – 50,801 | 21,033 | 85.1 |
| Medium | 15 | 15,573 | 8,538 | 7,247 – 35,767 | 17,293 | 88.5 |
| High | 15 | 10,191 | 6,300 | 3,284 – 24,780 | 11,357 | 87.5 |
| Top | 15 | 2,693 | 1,416 | 842 – 4,974 | 2,826 | 94.5 |
| Cumulative fall peak (CFP) |  |  |  |  |  |  |
| Site | N | CFP Mean | CFP StdDev | CFP Range | CND2 | CFP (%) |
| Low | 15 | 2,887 | 2,456 | 507 – 9,462 | 21,033 | 14.9 |
| Medium | 15 | 1,720 | 1,303 | 308 – 4,361 | 17,293 | 11.5 |
| High | 15 | 1,166 | 925 | 249 – 2,763 | 11,357 | 12.5 |
| Top | 15 | 134 | 129 | 14 – 399 | 2,826 | 5.5 |

Under the direct development hypothesis, we predict that the fall peak in year *y*-1 will be strongly correlated with the spring peak in year *y*. In contrast, under the delayed diapause hypothesis, we predict that the fall peak in year *y* should be strongly correlated with the spring peak in year *y*. To compare these two competing hypotheses, we created scatter plots of the fall peak versus the spring peak with different time lags and calculated the Pearson correlation coefficient. The fall peak in year *y*-1 was strongly correlated with the spring peak in year *y* for the low elevation site (Figure 4; r = 0.866, p = 0.0001) and the medium elevation site (Figure 4; r = 0.730, p = 0.005). In contrast, the fall peak and the spring peak in the same calendar year were not correlated (Additional file 1: Section 4). These results support the direct development hypothesis and provide strong evidence that the fall peak and spring peak that bookend the same winter represent the same generation of ticks.

**Figure 4.**
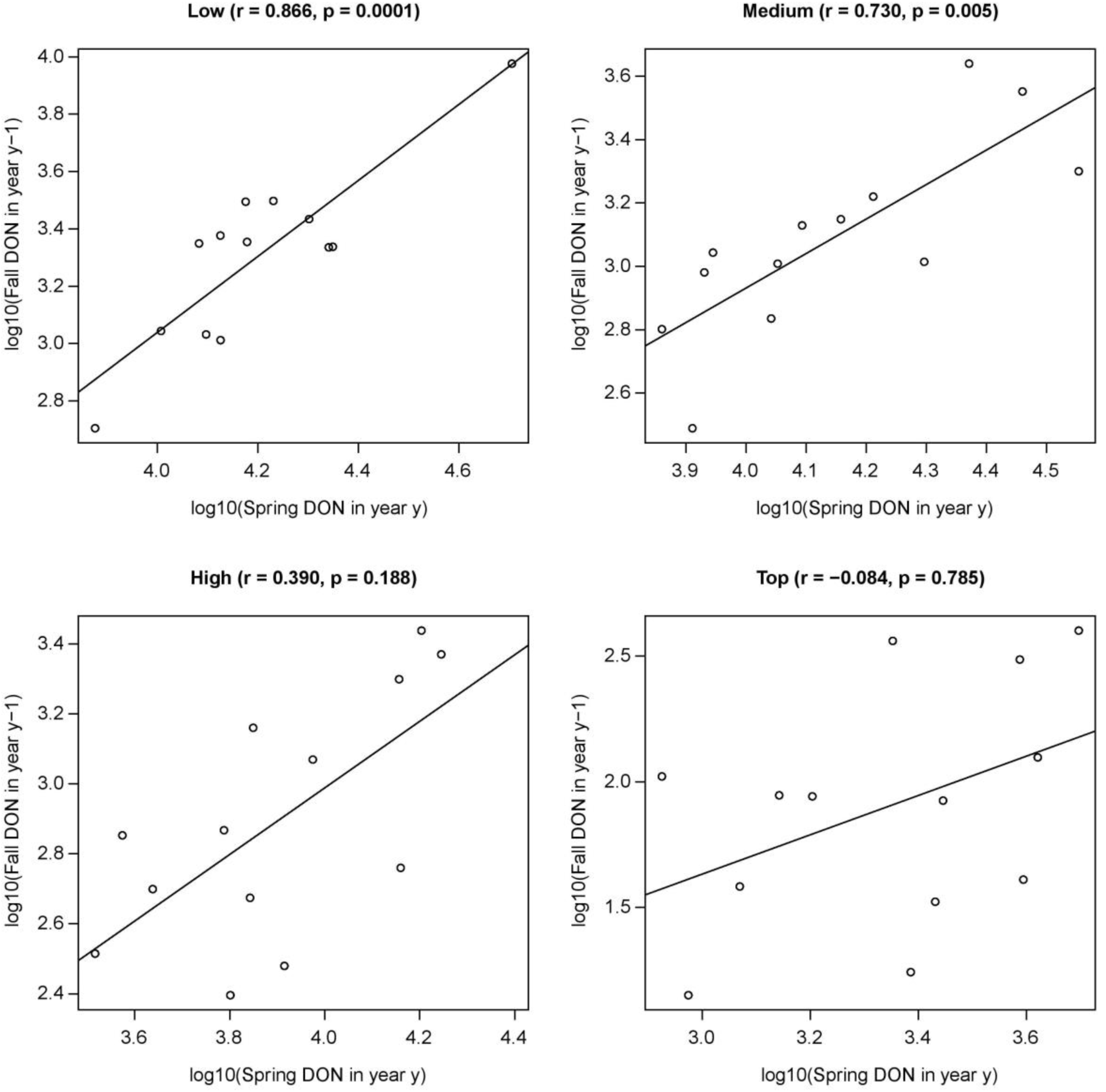
Correlation plot showing the relationship between the fall peak in year *y*-1 and the spring peak in year *y* for each of the four elevation sites. The fall peak in year *y*-1 is strongly correlated with the spring peak in year *y* for the low and medium elevation sites. The Pearson correlation coefficient (r) and the p-value (p) are shown in brackets at the top of each panel. These results support the direct development hypothesis and indicate that the tick year starts in the fall and ends the following summer.

We found that the bimodal non-linear phenology of the DON was best captured using GAMs that contained a non-parametric smoother function of the calendar day rather than temperature, relative humidity, or saturation deficit. Visual inspection shows that the values predicted by this smoother function of calendar day re-created the bimodal non-linear phenology of the DON (Additional file 1: Section 5).

### Sequential modelling approach

To determine the best model for explaining variation in the monthly DON over the 14 years of the study (2004 to 2017) at the four elevation sites on Chaumont Mountain, we used a sequential modelling approach (Additional file 1: Section 6). We used our previous work on the same data set as starting point [12, 36]. We compared more than 639 GAMs that modelled the DON as a function of the various explanatory variables using AIC-based model selection (Additional file 1: Section 6). Our sequential modelling approach led us to a set of 13 models that are presented in Table 5.

**Table 5.**
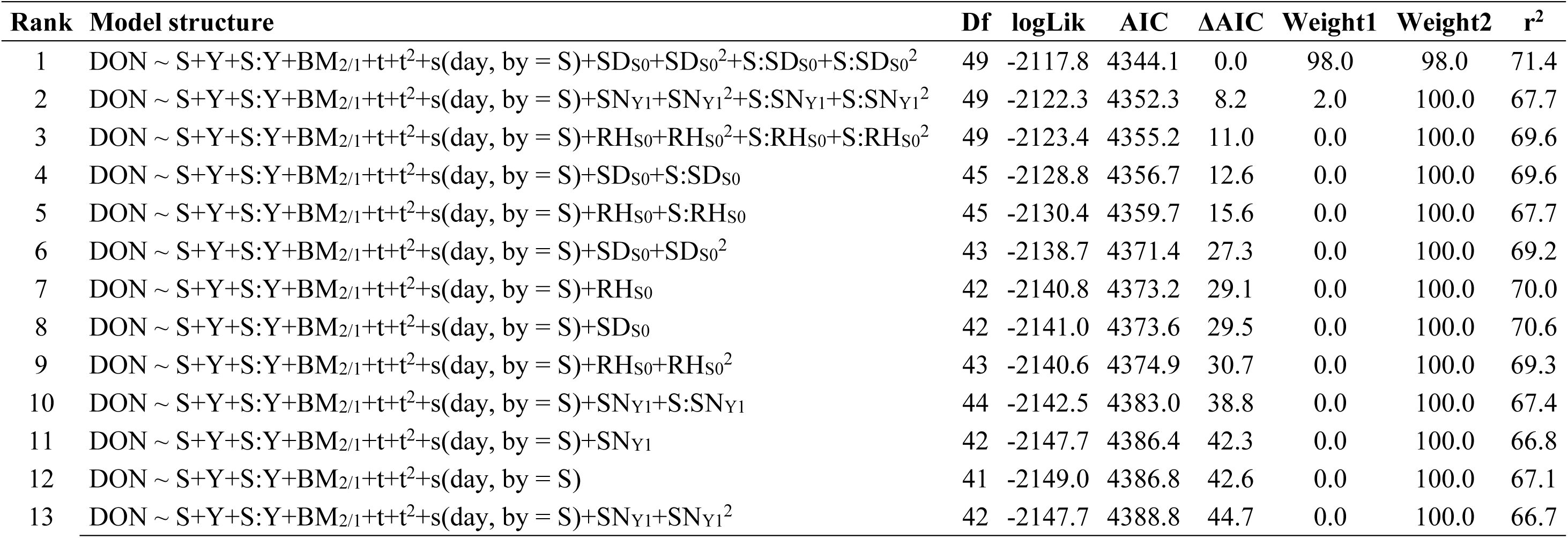
Model selection results are shown for the generalized additive model (GAM) with negative binomial errors of the density of *I. ricinus* nymphs (DON) at the four elevation sites on Chaumont Mountain over 14 years (2004 to 2017). *Notes:* The explanatory variables were site (S), year (Y), site:year interaction (S:Y), day (s(day, by = S)), beech mast score with different time lags for the spring peak and fall peak (BM_2/1_), temperature on day of sampling (t+t^2^), and 3 important weather station climate variables (SD_S0_, SN_Y1_, and RH_S0_). All climate variables were modelled as linear or quadratic effects. To model the bimodal non-linear phenology of the DON, a site-specific smoother function was applied to the calendar day, s(day, by = S). The models are ranked according to their Akaike information criterion (AIC). Shown for each model are the model rank (Rank), model structure (see Table 2 for the list of acronyms of the explanatory variables), model degrees of freedom (Df), log-likelihood (logLik), Akaike information criterion (AIC), difference in the AIC value from the top model (ΔAIC), model weight (Weight1), cumulative model weight (Weight2), and adjusted r-squared value (r^2^).

| Rank | Model structure | Df | logLik | AIC | ΔAIC | Weight1 | Weight2 | r<sup>2</sup> |
| --- | --- | --- | --- | --- | --- | --- | --- | --- |
| 1 | DON ~ S+Y+S:Y+BM<sub>2/1</sub>+t+t<sup>2</sup>+s(day, by = S)+SD<sub>S0</sub>+SD<sub>S0</sub><sup>2</sup>+S:SD<sub>S0</sub>+S:SD<sub>S0</sub><sup>2</sup> | 49 | -2117.8 | 4344.1 | 0.0 | 98.0 | 98.0 | 71.4 |
| 2 | DON ~ S+Y+S:Y+BM<sub>2/1</sub>+t+t<sup>2</sup>+s(day, by = S)+SN<sub>Y1</sub>+SN<sub>Y1</sub><sup>2</sup>+S:SN<sub>Y1</sub>+S:SN<sub>Y1</sub><sup>2</sup> | 49 | -2122.3 | 4352.3 | 8.2 | 2.0 | 100.0 | 67.7 |
| 3 | DON ~ S+Y+S:Y+BM<sub>2/1</sub>+t+t<sup>2</sup>+s(day, by = S)+RH<sub>S0</sub>+RH<sub>S0</sub><sup>2</sup>+S:RH<sub>S0</sub>+S:RH<sub>S0</sub><sup>2</sup> | 49 | -2123.4 | 4355.2 | 11.0 | 0.0 | 100.0 | 69.6 |
| 4 | DON ~ S+Y+S:Y+BM<sub>2/1</sub>+t+t<sup>2</sup>+s(day, by = S)+SD<sub>S0</sub>+S:SD<sub>S0</sub> | 45 | -2128.8 | 4356.7 | 12.6 | 0.0 | 100.0 | 69.6 |
| 5 | DON ~ S+Y+S:Y+BM<sub>2/1</sub>+t+t<sup>2</sup>+s(day, by = S)+RH<sub>S0</sub>+S:RH<sub>S0</sub> | 45 | -2130.4 | 4359.7 | 15.6 | 0.0 | 100.0 | 67.7 |
| 6 | DON ~ S+Y+S:Y+BM<sub>2/1</sub>+t+t<sup>2</sup>+s(day, by = S)+SD<sub>S0</sub>+SD<sub>S0</sub><sup>2</sup> | 43 | -2138.7 | 4371.4 | 27.3 | 0.0 | 100.0 | 69.2 |
| 7 | DON ~ S+Y+S:Y+BM<sub>2/1</sub>+t+t<sup>2</sup>+s(day, by = S)+RH<sub>S0</sub> | 42 | -2140.8 | 4373.2 | 29.1 | 0.0 | 100.0 | 70.0 |
| 8 | DON ~ S+Y+S:Y+BM<sub>2/1</sub>+t+t<sup>2</sup>+s(day, by = S)+SD<sub>S0</sub> | 42 | -2141.0 | 4373.6 | 29.5 | 0.0 | 100.0 | 70.6 |
| 9 | DON ~ S+Y+S:Y+BM<sub>2/1</sub>+t+t<sup>2</sup>+s(day, by = S)+RH<sub>S0</sub>+RH<sub>S0</sub><sup>2</sup> | 43 | -2140.6 | 4374.9 | 30.7 | 0.0 | 100.0 | 69.3 |
| 10 | DON ~ S+Y+S:Y+BM<sub>2/1</sub>+t+t<sup>2</sup>+s(day, by = S)+SN<sub>Y1</sub>+S:SN<sub>Y1</sub> | 44 | -2142.5 | 4383.0 | 38.8 | 0.0 | 100.0 | 67.4 |
| 11 | DON ~ S+Y+S:Y+BM<sub>2/1</sub>+t+t<sup>2</sup>+s(day, by = S)+SN<sub>Y1</sub> | 42 | -2147.7 | 4386.4 | 42.3 | 0.0 | 100.0 | 66.8 |
| 12 | DON ~ S+Y+S:Y+BM<sub>2/1</sub>+t+t<sup>2</sup>+s(day, by = S) | 41 | -2149.0 | 4386.8 | 42.6 | 0.0 | 100.0 | 67.1 |
| 13 | DON ~ S+Y+S:Y+BM<sub>2/1</sub>+t+t<sup>2</sup>+s(day, by = S)+SN<sub>Y1</sub>+SN<sub>Y1</sub><sup>2</sup> | 42 | -2147.7 | 4388.8 | 44.7 | 0.0 | 100.0 | 66.7 |

### Inter-annual variation in the fall and spring peaks of the DON are explained by different masting years

To investigate which of the 3 beech masting variables, BM_2/2_, BM_1/1_, BM_2/1_, was best at explaining the inter-annual variation in the spring and fall nymphal peaks, we compared their performance on a common background model (i.e., model 1 in Table 5). The best model contained the masting explanatory variable BM_2/1_, and had an AIC score of 4344.1, a support of 100.0%, and an r^2^ value of 71.4%. In contrast, the models containing BM_2/2_ and BM_1/1_ had AIC scores of 4481.3 and 4510.4 (ΔAIC with BM_2/1_ was 137.2 and 166.3), no support, and r^2^ values of 64.6% and 60.3%, respectively (Additional file 1: Section 7). A comparison of the model weights found that the support for the model with BM_2/1_ was nonillion times greater (1*10^30) compared to the model with BM_2/2_. Thus, the inter-annual variation in the spring and fall nymphal peaks are best explained by masting events that occurred 2 years prior and 1 year prior, respectively, which suggests that these two populations of nymphs are from different cohorts (i.e., born in different calendar years). This result provides strong evidence for the direct development hypothesis of the origin of the fall peak of nymphs.

### AIC-based model selection analysis of the best model

The model selection table for all 13 models is presented in Table 5. For the monthly DON, the best model had a support of 98.0%, explained 71.4% of the variation in the DON, and contained the explanatory variables of site (partial r^2^ = 13.8%), year (partial r^2^ = 6.6%), site:year interaction (partial r^2^ = 7.6%), beech mast score with different time lags for spring and fall peak (BM_2/1_; partial r^2^ = 16.0%), quadratic function of the temperature on the day of tick sampling (t and t^2^; partial r^2^ = 1.4%), quadratic function of the weather station mean seasonal SD of the summer in the present year (SD_S0_, SD_S0_, site:SD_S0_, and site:SD_S0_; partial r = 4.3%), and the smoothed function of the calendar day for each of the four elevation sites (partial r^2^ = 32.1%; Table 5).

The support for the 11 most important individual explanatory variables is shown in Table 6 and was as follows: site (100.0%), year (100.0%), site:year interaction (100.0%), beech mast score with different time lags for spring and fall peak (BM_2/1_; 100.0%), temperature on the day of tick sampling (support is 100.0% for both t and t^2^), smoothed function of the calendar day (100.0%), weather station mean seasonal SD of the summer in the present year (support is 97.8% to 97.9% for SD_S0_, SD_S0_, site:SD_S0_ interaction, and the site:SD_S0_ interaction). None of the other explanatory variables had a support > 2.0% (Additional file 1: Section 8).

**Table 6.**
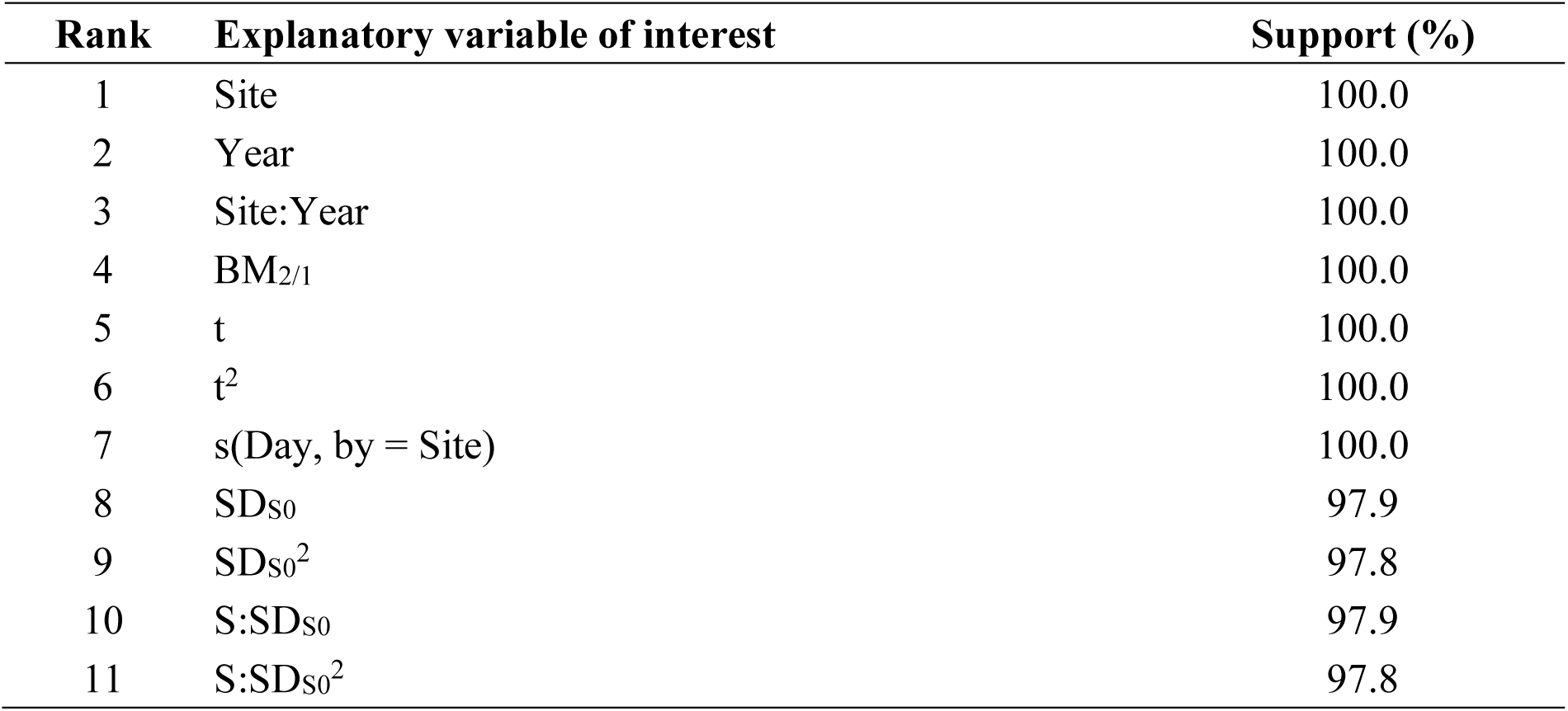
Support for the 11 most important explanatory variables of the GAMs of the DON. Notes: The support for the 11 most important explanatory variables is shown from the AIC-based model selection table of the GAMs of the DON. This support is calculated as the sum of the Akaike weights for all the models in the set that include that particular explanatory variable. Additional file 1: Section 7 shows the results for all the explanatory variables.

### Parameter estimates for the explanatory variables

To determine the effect of the explanatory variables on the DON, we present the parameter estimates on the log scale (and their 95% confidence intervals; Table 7). We also back calculated the effect sizes of the explanatory variables on the DON on the original scale with respect to the following reference conditions: the site was low elevation, the year was 2004, and the beech tree mast index was 1. For simplicity, we present the parameter estimates from the top model (model 1 in Table 5), but we note that they are similar to the model-averaged parameter estimates (Additional file 1: Section 8).

**Table 7.**
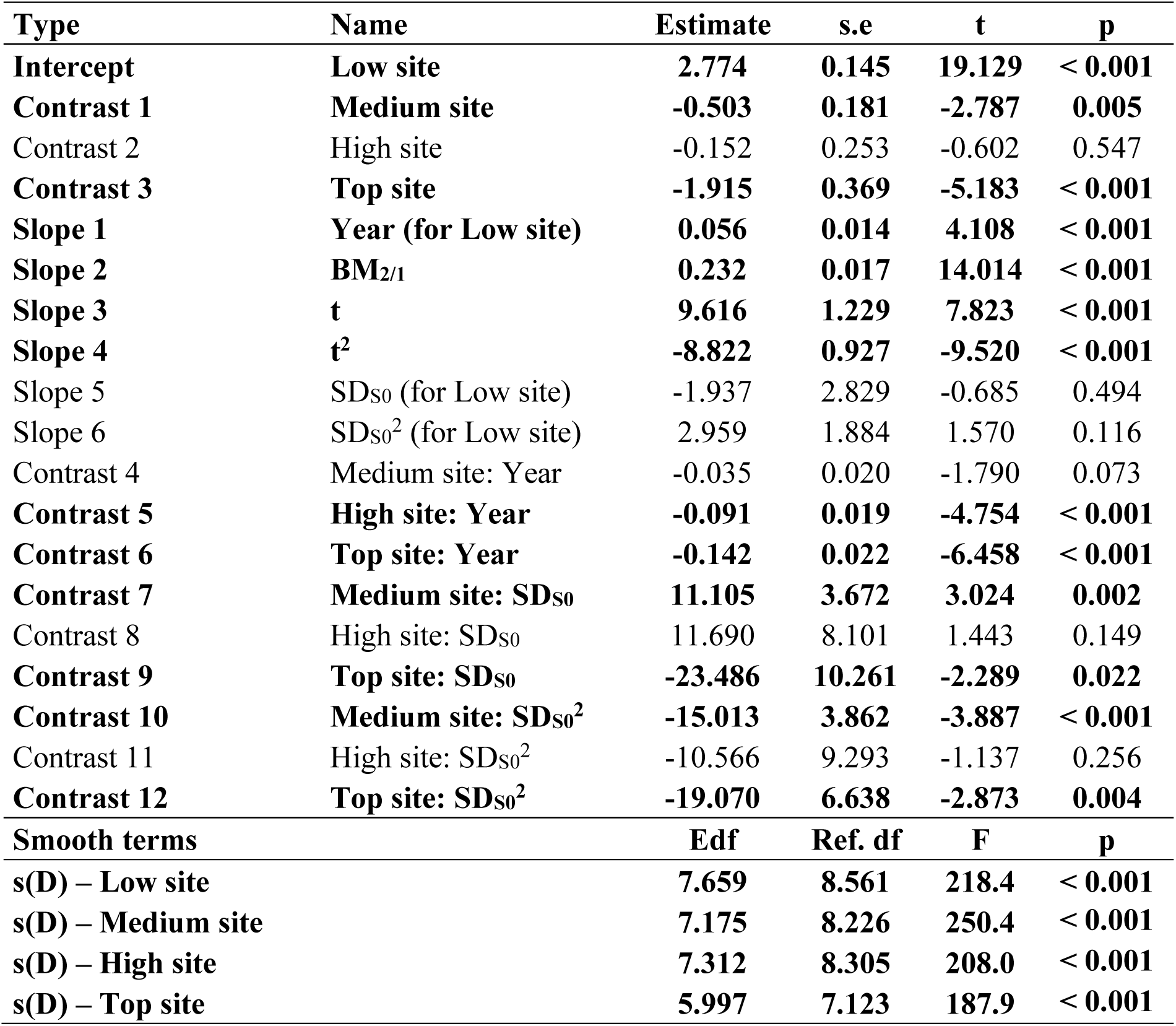
The parameter estimates from the best model in Table 5 are shown. *Notes:* In this best model, the DON response variable was modelled as a function of elevation site (Low, Medium, High, and Top), year, beech mast score with different time lags for the spring peak and fall peak (BM_2/1_), linear and quadratic terms for temperature on the day of tick sampling (t and t^2^), linear and quadratic terms for the weather station mean seasonal saturation deficit of the summer in the present year (SD_S0_ and SD_S0_). Interactions are indicated by colons (:); for example, the parameter estimates for the site:year interaction are indicated by the terms Medium site: Year, High site: Year, and Top site: Year. For each parameter, the parameter type, parameter name, parameter estimate on the log scale, standard error on the log scale (s.e.), t-statistic (t), and p-values (p) are shown. To model the bimodal non-linear phenology of the DON, a site-specific smoother function was applied to the calendar day. For each of the 4 site-specific smoother functions of the calendar day, the effective degrees of freedom (edf), reference degrees of freedom (Ref. df), F-statistic (F), and p-value (p) are shown.

| Type | Name | Estimate | s.e | t | p |
| --- | --- | --- | --- | --- | --- |
| **Intercept** | **Low site** | **2.774** | **0.145** | **19.129** | **< 0.001** |
| **Contrast 1** | **Medium site** | **-0.503** | **0.181** | **-2.787** | **0.005** |
| Contrast 2 | High site | -0.152 | 0.253 | -0.602 | 0.547 |
| **Contrast 3** | **Top site** | **-1.915** | **0.369** | **-5.183** | **< 0.001** |
| **Slope 1** | **Year (for Low site)** | **0.056** | **0.014** | **4.108** | **< 0.001** |
| **Slope 2** | **BM<sub>2/1</sub>** | **0.232** | **0.017** | **14.014** | **< 0.001** |
| **Slope 3** | **t** | **9.616** | **1.229** | **7.823** | **< 0.001** |
| **Slope 4** | **t<sup>2</sup>** | **-8.822** | **0.927** | **-9.520** | **< 0.001** |
| Slope 5 | SD<sub>s0</sub> (for Low site) | -1.937 | 2.829 | -0.685 | 0.494 |
| Slope 6 | SD<sub>s0</sub><sup>2</sup> (for Low site) | 2.959 | 1.884 | 1.570 | 0.116 |
| Contrast 4 | Medium site: Year | -0.035 | 0.020 | -1.790 | 0.073 |
| **Contrast 5** | **High site: Year** | **-0.091** | **0.019** | **-4.754** | **< 0.001** |
| **Contrast 6** | **Top site: Year** | **-0.142** | **0.022** | **-6.458** | **< 0.001** |
| **Contrast 7** | **Medium site: SD<sub>s0</sub>** | **11.105** | **3.672** | **3.024** | **0.002** |
| Contrast 8 | High site: SD<sub>s0</sub> | 11.690 | 8.101 | 1.443 | 0.149 |
| **Contrast 9** | **Top site: SD<sub>s0</sub>** | **-23.486** | **10.261** | **-2.289** | **0.022** |
| **Contrast 10** | **Medium site: SD<sub>s0</sub><sup>2</sup>** | **-15.013** | **3.862** | **-3.887** | **< 0.001** |
| Contrast 11 | High site: SD<sub>s0</sub><sup>2</sup> | -10.566 | 9.293 | -1.137 | 0.256 |
| **Contrast 12** | **Top site: SD<sub>s0</sub><sup>2</sup>** | **-19.070** | **6.638** | **-2.873** | **0.004** |
| **Smooth terms** |  | **Edf** | **Ref. df** | **F** | **p** |
| **s(D) – Low site** |  | **7.659** | **8.561** | **218.4** | **< 0.001** |
| **s(D) – Medium site** |  | **7.175** | **8.226** | **250.4** | **< 0.001** |
| **s(D) – High site** |  | **7.312** | **8.305** | **208.0** | **< 0.001** |
| **s(D) – Top site** |  | **5.997** | **7.123** | **187.9** | **< 0.001** |

The interaction between elevation site and year indicated that the change in the DON over time differed between the four elevation sites (Figure 5; Table 7). Over the 14-year period (2004–2017), the DON increased at the low elevation (slope = 0.056 per year, s.e. = 0.014, p < 0.001), and medium elevation (Medium – Low contrast of the slope = −0.035, s.e. = 0.020, p = 0.073) sites, but decreased at the high elevation (High – Low contrast of the slope = −0.091, s.e. = 0.019, p < 0.001), and top elevation (Top – Low contrast of the slope = −0.142, s.e. = 0.022, p < 0.001) sites. Over the 14-year period (2004–2017), the DON increased by 119.0% and 34.2% at the low and medium elevation sites but decreased by 38.7% and 70.0% at the high and top elevation sites, respectively (Figure 5).

**Figure 5.**
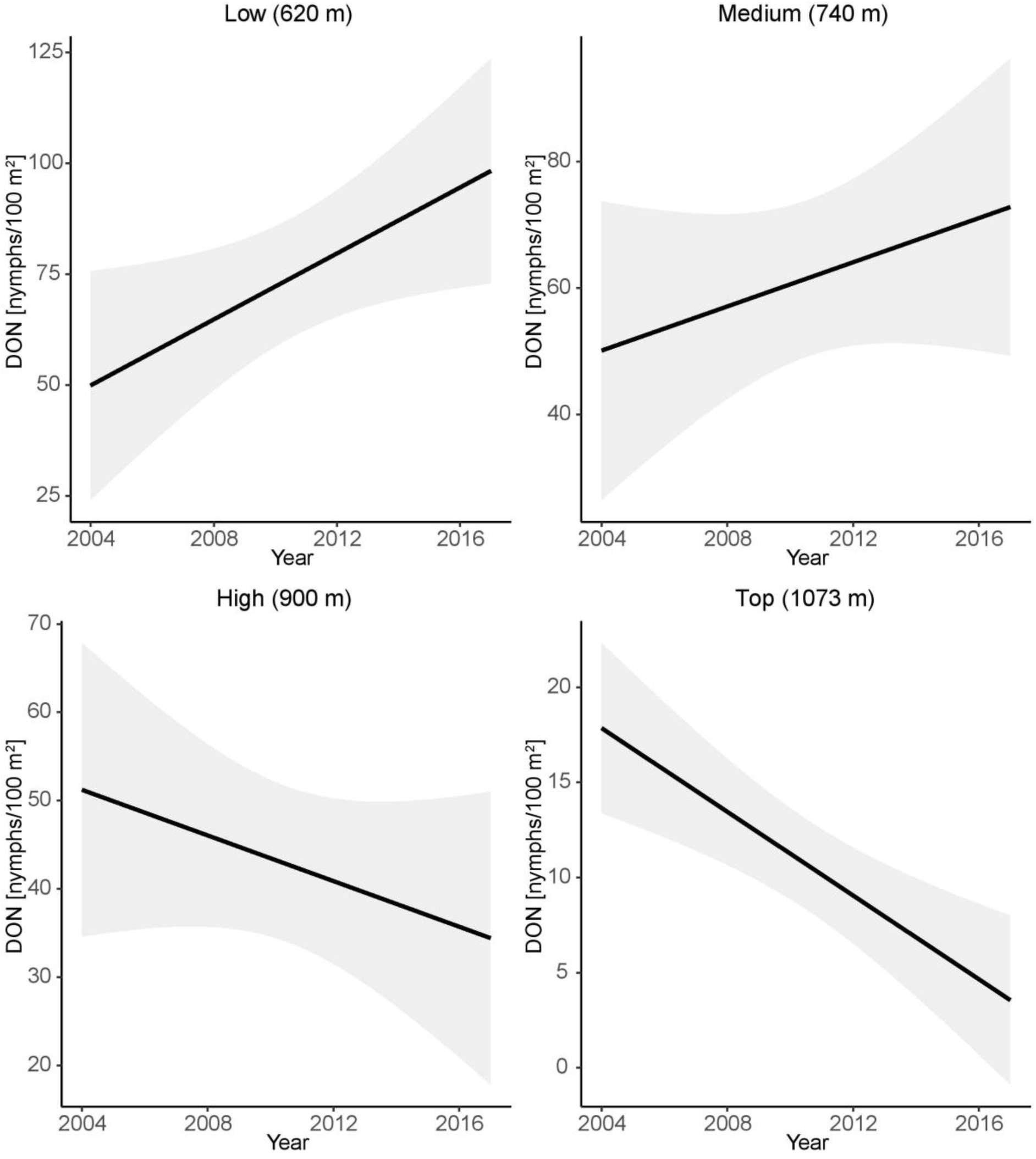
Effect of year on the density of nymphs (DON). The DON is an estimate of the number of questing *I. ricinus* nymphs per 100 m^2^ sampled by the dragging method each month. The parameter estimates used to calculate the effect sizes were taken from Table 7. Over the 14-year study period, the DON increased by 119.0% at the low elevation, increased by 34.2% at the medium elevation, decreased by 38.7% at the high elevation, and decreased by 70.0% at the top elevation (partial r^2^ = 6.6%).

The beech mast score with different time lags (2 years versus 1 year) for the spring and fall peak (BM_2/1_) had a positive and significant effect on the DON (slope = 0.232 per class, s.e. = 0.017, p < 0.001; Figure 6; Table 7). Increasing the beech mast score from 1 (poor mast) to 5 (full mast) increased the DON by 152.9% at each of the four elevation sites on Chaumont Mountain (Figure 6).

**Figure 6.**
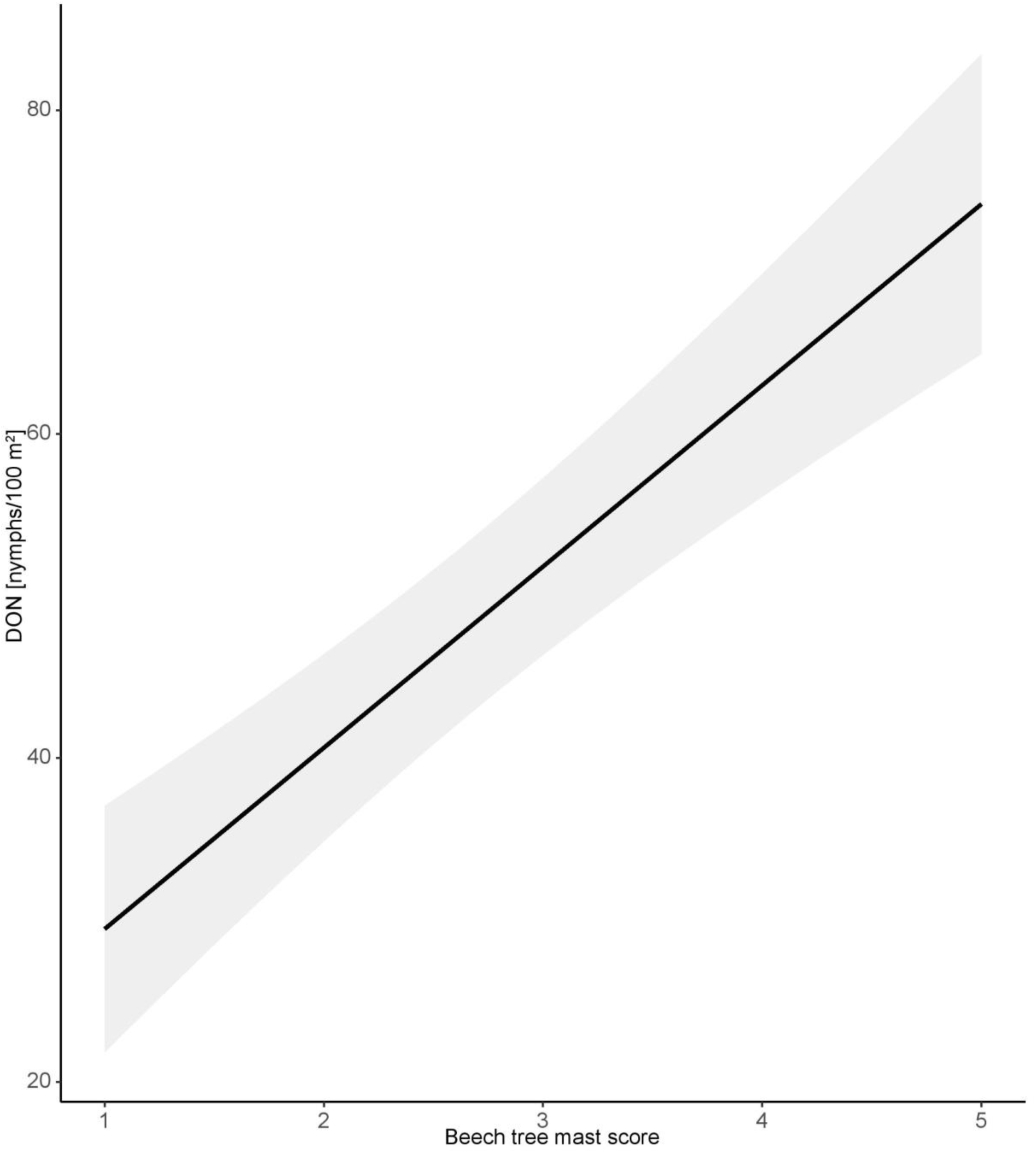
Effect of beech mast score with different time lags (2 years versus 1 year) for the spring and fall peak (BM_2/1_) on the density of nymphs (DON). The beech tree mast score (BM_2/1_) assumes a 2-year time lag for the spring nymphal peak and a 1-year time lag for the fall nymphal peak. The linear relationship shows that the DON increases with the beech mast score. The DON is an estimate of the number of questing *I. ricinus* nymphs per 100 m^2^ sampled by the dragging method each month. Beech tree mast scores have values of 1, 2, 3, 4, and 5, which refer to very poor mast, poor mast, moderate mast, good mast, and full mast, respectively. The parameter estimates used to calculate the effect sizes were taken from Table 7. Increasing the beech mast score from 1 (poor mast) to 5 (full mast) increased the DON by 152.9% at each of the four elevation sites (partial r^2^ = 16.0%).

The slope of the linear effect of the field-measured temperature (i.e., the temperature measured on the day of tick sampling) on the DON was positive and significant (slope = 9.616 per °C, s.e. = 1.229, p < 0.001; Figure 7; Table 7), indicating that the DON increased with temperature over the range of observed values (−5°C to 30°C). The slope of the quadratic effect of the field-measured temperature on the DON was negative and significant (slope = −8.822, s.e. = 0.927, p < 0.001; Figure 7; Table 7) indicating that the DON plateaued once the field-measured temperature reached ∼ 30°C.

**Figure 7.**
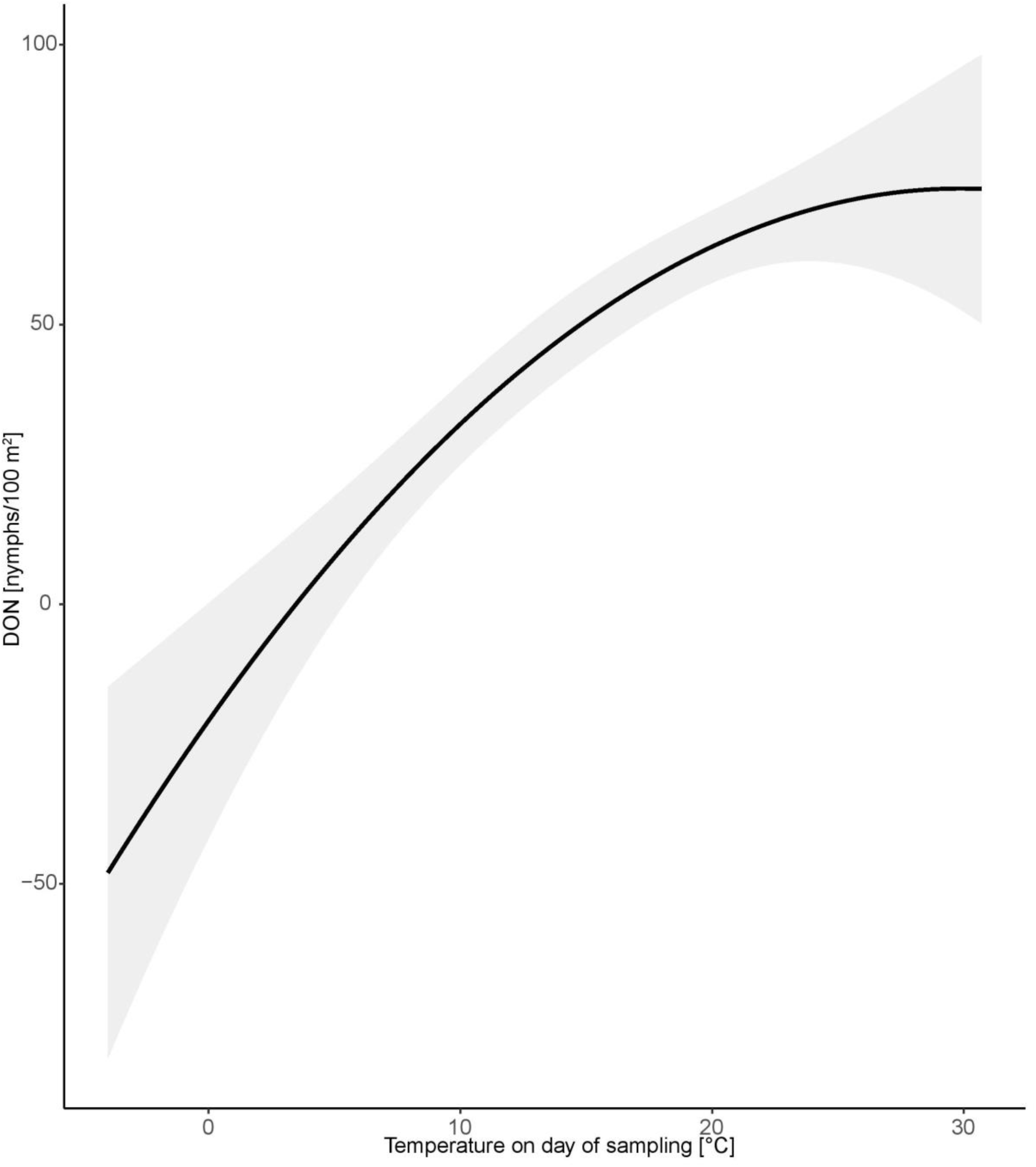
Effect of the field-measured temperature on the day of tick sampling on the density of nymphs (DON). The quadratic relationship shows that the DON increases with the temperature on the day of tick sampling and reaches a plateau at 30°C. The DON is an estimate of the number of questing *I. ricinus* nymphs per 100 m^2^ sampled by the dragging method each month. Temperature has units of °C and was measured at 60 cm above the ground at the field site on the day of tick sampling. The parameter estimates used to calculate the effect sizes were taken from Table 7. The quadratic function of the field-measured temperature explained 0.0% of the variation in the DON (partial r^2^ = 1.4%).

The SD_S0_ is the mean SD during the summer (1 June to 31 August) of the same year as the DON (i.e., no time lag). Thus, the SD_S0_ represents the SD during the last 3 months of the spring nymphal peak, just before the start of the fall peak. The significant interaction between elevation site and SD_S0_ indicates that the relationship between the DON and the SD_S0_ differed between the four elevation sites (Figure 8; Table 7). The relationship between the DON and the SD_S0_ was positive linear at the low and high elevation sites, and negative quadratic at the medium and top elevation sites. At the medium and top elevation sites, the DON decreased once the SD_S0_ reached ∼ 5.5 mmHg and ∼ 4.0 mmHg, respectively.

**Figure 8.**
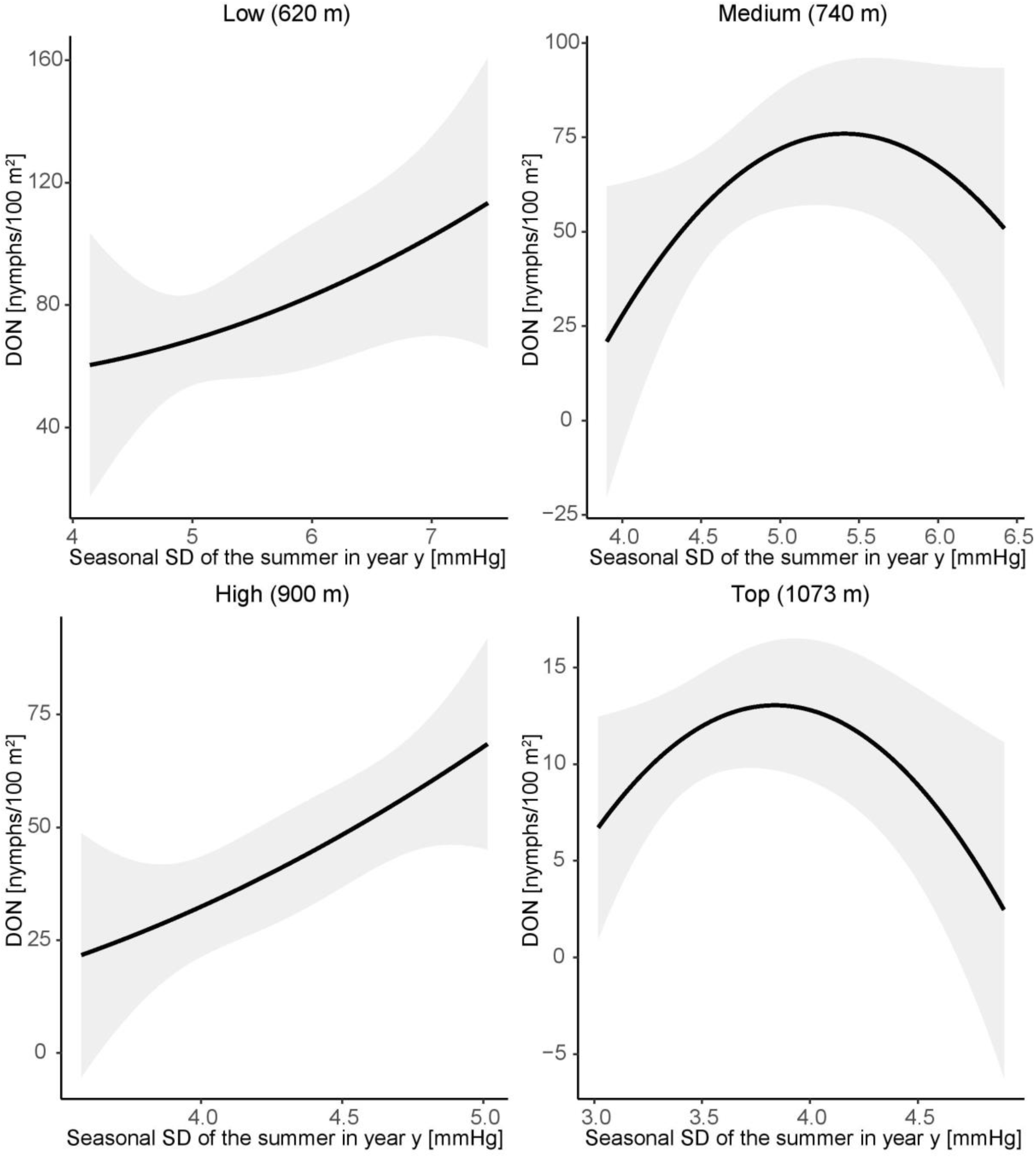
Effect of the mean saturation deficit in the summer of year *y* (SD_S0_) on the density of nymphs (DON) at each of the four elevation sites. The SD_S0_ refers to the mean saturation deficit calculated over the summer months (1 June to 31 August) in the same year as the DON (i.e., no time lag). The relationship between the DON and the SD_S0_ is linear and positive for the low and high elevation sites and it is a negative quadratic for the medium and top elevation sites. The DON is an estimate of the number of questing *I. ricinus* nymphs per 100 m^2^ sampled by the dragging method each month. The SD_S0_ has units of mmHg and was calculated from temperature and relative humidity data measured at 200 cm above the ground by two weather stations near the field site. The parameter estimates used to calculate the effect sizes were taken from Table 7. The site-specific quadratic function of SD_S0_ explained 14.3% of the variation in the DON (partial r^2^ = 4.3%).

In summary, the DON had a bimodal phenology at the low, medium, and high elevation sites and a unimodal phenology at the top elevation site. The DON increased over time at the low and medium elevation sites, whereas it decreased at the high and top elevation sites. The DON increased significantly with beech masting index and the time lag differed between the two nymphal peaks with a 2-year time lag for the spring peak and a 1-year time lag for the fall peak. This result provides strong evidence that the spring and fall nymphal peaks are distinct cohorts that are born in different years, which supports the direct development hypothesis and not the developmental diapause hypothesis. The DON increased with the field-measured temperature but reached a plateau at 30°C. Finally, the relationship between the DON and the seasonal SD of the summer in year *y* (SD_S0_) was positive linear at the low and high elevation sites, and negative quadratic at the medium and top elevation sites.

### Fall nymphs have higher fat content than the spring nymphs

We compared the log10-transformed fat content between the *I. ricinus* nymphs collected in the fall of 2010 and the nymphs collected in the spring of 2010 using a two independent samples t-test. The mean fat content of the fall nymphs (n = 40; 9.3 ug; 95% CI = 7.0 to 12.4) was 76.1% higher than that of the spring nymphs (n = 40; 5.3 ug; 95% CI = 4.0 to 7.1), and this difference was significant (Figure 9; t = 2.653, df = 68.003, p = 0.010). This result suggests that the fall nymphs are younger than the spring nymphs, which is consistent with the direct development hypothesis.

**Figure 9.**
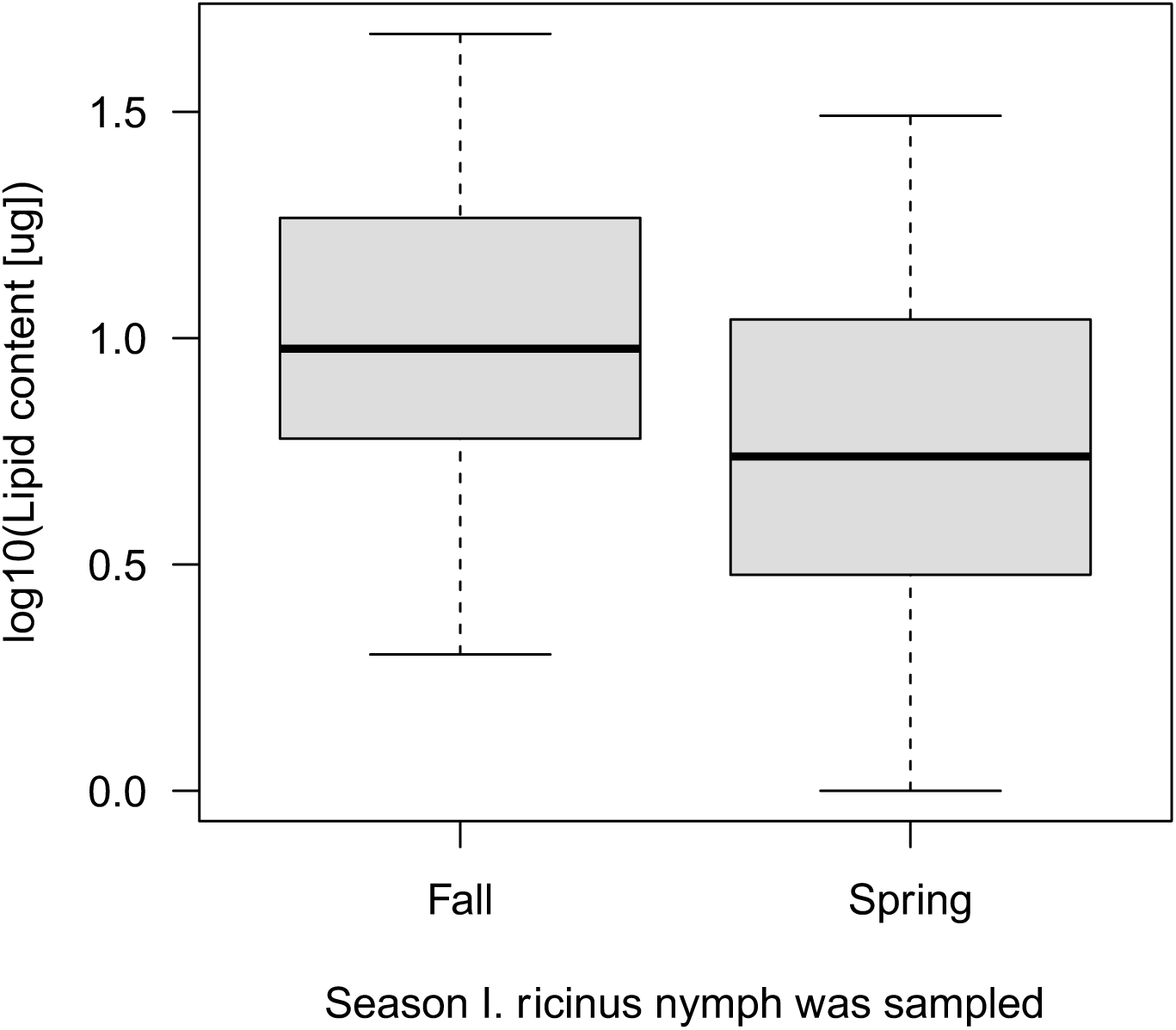
*I. ricinus* nymphs collected in the fall have a higher fat content compared to nymphs collected in the spring of the same calendar year. This result suggests that the fall nymphs are younger than the spring nymphs. The boxplot shows the medians (black line), the 25th and 75th percentiles (edges of the box), and the minimum and maximum values (whiskers).

## DISCUSSION

Given the importance of *I. ricinus* as a disease vector, forecasting the density of ticks questing for hosts is important for managing the risk of tick-borne diseases [12, 13, 35, 36, 69–71]. In Europe, there is much interest to determine which ecological factors are influencing the seasonal and inter-annual abundance of *I. ricinus* ticks [12, 35]. In continental Europe, *I. ricinus* nymphs have a bimodal seasonal phenology with a large spring peak followed by a smaller fall peak. There are at least two alternative hypotheses to explain the fall peak of nymphs: direct development and delayed diapause. We provide multiple lines of evidence that the fall peak is best explained by the direct development hypothesis at our field site in Switzerland. The fall peak of nymphs does not occur at our top elevation site because summer temperatures do not permit direct development of engorged larvae into nymphs. The fall peak in year *y*-1 is strongly correlated with the spring peak in year *y*, indicating that they represent the same tick generation. Inter-annual variation in the fall peak is best explained by a 1-year time lag with beech masting compared to the standard 2-year time lag for the spring peak. Fall nymphs have higher fat content than spring nymphs indicating that they are younger. All of these results are consistent with the direct development hypothesis and they are inconsistent with the delayed diapause hypothesis, as we explain in more detail below.

### The *I. ricinus* nymphs at Chaumont Mountain have a bimodal phenology

The seasonal phenology of *I. ricinus* nymphs at Chaumont Mountain is bimodal where a large spring peak is followed by a smaller fall peak. This bimodal phenology occurred at the three lower elevation sites, whereas a unimodal phenology with a single large peak in the spring occurred at the top elevation site (Figure 3), and this discrepancy is consistent with the direct development hypothesis. The mean field-measured temperature in the spring and summer at the top elevation site is 5.4°C colder compared to the low elevation site. The result is that at the top site, tick development rates are slower, engorged larvae rarely (if ever) undergo direct development to questing nymphs in the same calendar year, and there is no fall peak of nymphs. Across Europe, we expect *I. ricinus* populations to have bimodal phenologies where longer, warmer summers facilitate direct development of engorged larvae into unfed nymphs in the summer so that they have enough time to quest and create the fall nymphal peak that same year (**Table 1**). In contrast, we expect unimodal phenologies to occur at northern latitudes or at high elevation sites (**Table 1**) where lower temperatures during the summer do not allow engorged larvae to complete their development from engorged larvae to unfed nymphs and quest in the same calendar year [40, 45, 72].

Our study also found that *I. ricinus* nymphs started questing earlier at the low elevation site (April) compared to the medium, high, and top elevation sites (May), and this result is probably also true for the larvae. Thus, larvae at the low site start questing earlier, which means that they have more time to complete their larvae-to-nymph moult and at faster development rates due to the elevated summer temperatures compared to larvae at the top elevation site. All these differences combine to create a fall nymphal peak that is prominent at the low elevation site and absent at the top elevation site, and this altitudinal gradient in phenology is consistent with the direct development hypothesis.

### One cohort of larvae occurs in two different calendar years

A striking result of our study is the strong correlation between the fall nymphal peak in year *y*-1 and the spring nymphal peak in year *y* at both the low and medium elevation sites (Figure 4). In contrast, there is no correlation between the spring peak and the fall peak in the same calendar year (**Additional file 1: Section 4**). These results are consistent with the direct development hypothesis and they are not consistent with the delayed diapause hypothesis (Figure 1). Under the direct development hypothesis, the larvae that obtain an early larval blood meal and/or that develop fast in year *y*-1, will quest as nymphs in the fall of year *y*-1, whereas the larvae that obtain a late larval blood meal and/or that develop slow, will overwinter as unfed nymphs and quest in the spring of year *y.* Under the direct development hypothesis, the tick year starts in the fall of year *y*-1 and ends in the summer of year *y* and it does not correspond to the calendar year. Studies that analyze inter-annual variation in the density of *I. ricinus* typically calculate a cumulative DON for the calendar year [12, 35, 36, 44, 49]. The present study shows that this approach is wrong when the fall peak and the spring peak in the same calendar year are two different cohorts of ticks. Researchers studying *I. ricinus* populations with a bimodal phenology should be aware that the fall nymphal peak and the spring nymphal peak that bookend the winter may be part of the same cohort of nymphs.

### Beech masting has different time lags with the spring and fall nymphs

An important result is the strong and positive association between the beech masting score (BM_2/1_) and the density of *I. ricinus* nymphs in the spring (Figure 6). For the spring peak, the consensus is that it consists of larvae that obtained their blood meal the previous spring and summer, molted into nymphs in the summer and early fall (Kahl, 1989), entered behavioral diapause, overwintered as unfed nymphs, and quested the following spring [19, 45, 46]. We expect a 2-year time lag between beech masting and the spring nymphal peak because masting in the fall of year *y*-2 increases the density of mice and larval feeding success in the spring and summer of year *y*-1, which in turn increases the DON in the spring of year *y*. Studies in North America and Europe have shown that the masting events of deciduous trees can drive inter-annual variation in the DON and DIN of *Ixodes* nymphs with a 2-year time lag [14, 32, 33, 35]. We had previously shown using the same data from the present study that the beech masting score 2 years prior was highly significantly associated with inter-annual variation in the DON and DIN [12, 36]. In these studies, we had incorrectly assumed that the spring peak and fall peak that bookend the summer and occur in the same calendar year represent the same cohort of nymphs [12, 36]. Our previous studies found a strong association between beech masting 2 years prior and the cumulative DON because the spring peak is typically 6 to 7 times larger than the fall peak. Nevertheless, our decision to combine the spring and fall peak of the same calendar year into a single estimate of the annual DON and DIN was incorrect [12, 36].

The novel aspect of the present study is that our analysis of the bimodal phenology of the DON shows that the spring peak and the fall peak of the same calendar year represent different cohorts of nymphs. Inter-annual variation in the spring peak and fall peak was best predicted by the beach masting index 2 years prior and 1 year prior, respectively. This result provides strong evidence for the direct development hypothesis [38, 42, 43, 45, 46] which predicts a 1-year time lag between beech masting and the fall nymphal peak; masting in year *y*-1 increases the density of mice and larval feeding success in year *y*, which in turn increases the DON in the fall of year *<u>y</u>*. In contrast, the developmental diapause hypothesis [19, 45, 46] predicts a 2-year time lag between beech masting and the fall nymphal peak; masting in year *y*-2 increases the density of mice and larval feeding success in year *y*-1, but larvae that obtain their blood meal in late summer have to overwinter as engorged larvae, moult into unfed nymphs the following summer, and quest in the fall of year *y*. A seminal review on diapause in *Ixodes* ticks presented both hypotheses, but appeared to favor the developmental diapause hypothesis over the direct development hypothesis [19]. In contrast, we found that the direct development hypothesis for the fall peak of *I. ricinus* nymphs had 100% support whereas the developmental diapause hypothesis had 0% support.

### Differences in fat content between spring and fall nymphs support the direct development hypothesis

The direct development hypothesis predicts that the fall nymphs are younger compared to the spring nymphs (∼3 months versus ∼9 months since the larval blood meal), which agrees with studies comparing the fat content between these two types of nymphs in our study area and elsewhere [45, 63]. In the present re-analysis of the fat content of field-collected *I. ricinus* nymphs at our study location (Figure 9), we found that the fall nymphs had 76% more fat content than the spring nymphs [63], and studies in the UK have found a similar pattern [45]. This phenomenon can be explained by the direct development hypothesis; fall nymphs obtained their larval blood meal earlier that summer and their fat content is high because they are young (∼3 months since the larval blood meal) with less time to burn their fat reserves. In contrast, spring nymphs obtained their larval meal the previous summer and their fat content is low because they are older (∼9 months since the larval blood meal) with more time to burn their fat reserves. Thus, the finding that fall nymphs have higher fat reserves than spring nymphs is consistent with our discovery of 1-year and 2-year time lags for the fall and spring peak, and both these results support the direct development hypothesis.

Some studies have suggested a third explanation for the fall peak of *I. ricinus* nymphs. The summer quiescence hypothesis proposes that nymphal questing activity decreases in the late summer because the nymphs are hiding in the leaf litter from unfavorable conditions (high temperatures and low RH) and waiting to resume their questing activity in the fall when conditions are more favorable [18, 42]. We disagree with this hypothesis for two reasons. First, if the spring and fall peak are from the same cohort then the fall nymphs are older and should have lower fat content than the spring nymphs, but this is not the case [45, 63]. Second, if the spring and fall nymphs are from the same cohort then they should have the same time lag with the masting index, but this was not the case in our study. In summary, our study does not support the summer quiescence hypothesis.

### Effect of weather on the questing behavior of *I. ricinus* nymphs

Our study found that the field-measured temperature on the day of tick sampling was positively correlated with the density of questing nymphs (Figure 7), which agrees with other studies in Europe [39, 73–76]. These studies suggest that for any given date, warmer temperatures increase the percentage of nymphs questing for hosts (i.e., increased nymphal activity levels), which in turn, increase the number of questing nymphs captured by drag sampling. *I. ricinus* becomes active above a minimum threshold temperature and questing activity increases with temperature up to some maximum value. While these threshold temperatures are expected to vary among tick populations from different geographical locations, a previous study at our field location suggested that the maximum threshold value was 24°C [42], whereas the present study found that this threshold was 30°C. *I. ricinus* is also sensitive to desiccation during questing [17], and they make repeated return trips to the litter layer where they can rehydrate and maintain their water balance [51, 52, 77]. For this reason, ticks prefer to quest under cool and humid conditions (i.e., low SD) [48, 78] and they reduce their questing activity in hot and dry conditions (i.e., high SD) [42, 50, 58, 79, 80]. Our observation that the DON plateaued at ∼30°C in our study (Figure 7) suggests that *I. ricinus* nymphs at our study site avoid questing at hot (and presumably dry) conditions.

### Effect of climate variables on the population ecology of *I. ricinus* ticks

Climate variables can influence tick population ecology via their effects on the vital rates (development, survival, reproduction). An interesting result of our study was that the seasonal climate variables were much more important than the annual climate variables at explaining the inter-annual variation in the DON. Our previous work found that annual climate variables (e.g., mean annual relative humidity and mean annual precipitation) were important for explaining the inter-annual variation in the DON or DIN [12, 36], but these studies did not investigate seasonal means. The present study is better because it compares climate variables operating at different temporal scales (i.e., annual versus seasonal). Many steps in the tick life cycle happen over short time scales suggesting that the critical climate variables operate over seasons rather than years. Researchers have addressed this issue by investigating a wide variety of durations and time lags [13, 49, 81]. For example, a 14-year study in Switzerland averaged the climate variables over different time intervals (1, 5, 10, 17 or 30 days) to determine the critical time period that influences the DON [49]. An 8-year study in Germany used cross correlation maps to explore month-to-month correlations between the DON and climate variables, as well as time-lagged and interval-averaged correlations by considering a second time lag [13]. A 2-year study in five European countries examined time-lagged and interval-aggregated monthly and yearly means to test the associations between the abundance of questing *I. ricinus* nymphs and climate [81]. Despite this research effort, there is still uncertainty about which climate variables operating over what time scales drive the population ecology of *I. ricinus*.

Our study found that the relationship between the mean saturation deficit in the summer of that same year (SD_S0_) and the DON differed among the four elevation sites (Figure 8). The relationship was positive linear for the low and high elevation sites, and it was negative quadratic for the medium and top elevation sites (Figure 8). The SD_S0_ is expected to have different effects on the spring and fall nymphs. With respect to the spring nymphs, increasing the SD is initially expected to increase the nymphal questing activity (the positive effects of higher temperatures outweigh the negative effects of lower relative humidity). However, the risk of desiccation and death both increase with high SD [50, 82], and at a certain threshold SD value, questing activity and survival will start to decrease for the spring nymphs (the negative effects of lower relative humidity outweigh the positive effects of higher temperatures). Thus, for the spring nymphs we expect a negative quadratic relationship between SD_S0_ and the DON, as observed at the medium and top elevation sites (Figure 8). With respect to the fall nymphs, a hot and dry summer (high SD) accelerates tick developmental rates, which under the direct development hypothesis, would increase the size of the fall nymphal peak of that calendar year [38, 42, 43, 45, 46]. In summary, the observed relationships between the SD_S0_ and the DON are broadly consistent with our expectations.

All three of our studies on this 15-year data set have found that RH (either acting alone or in combination with temperature to form the SD) in the present year (either annual or seasonal) was the most important climate variable for explaining variation in the DON or DIN [12, 36]. In our first study, the field-measured mean annual RH in the present year was negatively correlated with the annual DON [12]. In our second study, the weather station mean annual RH in the present year was negatively correlated with the DIN. In the present study, the best models in Table 5 included the summer SD in the present year (SD_S0_) and the summer RH in the present year (RH_S0_). However, one contradictory result in the present study was that the RH_S0_ was positively correlated with the DON (**Additional file 1: Section 8**). Thus, RH was always important for the ecology of *I. ricinus* but its correlation with the DON (negative or positive) differed among the three studies.

### GAMs can model the bimodal phenology of *I. ricinus* nymphs

The bimodal non-linear phenology of *I. ricinus* nymphs is a challenge for statistical modelling. In our previous work, we avoided this complexity by analysing the total DON and DIN for each year and the same approach was used in a 14-year study in Switzerland that is near our study site [49], in a 10-year study in the Netherlands [44], and in a 13-year study in North America [14]. An 8-year study in Germany used different intercepts for the spring, summer, fall, and winter to deal with the non-linear phenology of *I. ricinus* nymphs [13]. We demonstrated that generalized additive models (GAMs) are a useful approach to model the bimodal non-linear phenology of *I. ricinus* nymphs because they allow the user to flexibly model the response variable as a non-parametric and non-linear smoothed function of the explanatory variables of interest. The smoothed function of the calendar day was an excellent fit to the seasonal phenology of *I. ricinus* nymphs (**Additional file 1: Section 5**). The other explanatory variables (e.g., beech masting score and climate variables) were initially modelled with non-parametric smoothed functions because we did not want to assume a linear function, but were subsequently replaced with parametric functions (e.g., linear or quadratic). In summary, we demonstrated that the bimodal non-linear phenology of *I. ricinus* ticks can be captured by GAMs using a combination of parametric and non-parametric functions of the explanatory variables of interest.

### Conclusions

At our study location, the *I. ricinus* populations have a bimodal phenology at the lower elevation sites, but a unimodal phenology at the top elevation site. At the low and medium elevation sites, the fall nymphal peak in year *y*-1 was strongly correlated with the spring nymphal peak in year *y* over the 15 years of the study. In contrast, there was no correlation between the fall peak and the spring peak in the same calendar year. Inter-annual variation in the fall nymphal peak and spring nymphal peak were strongly associated with the beech masting index 1 year prior and 2 years prior, respectively. Fall nymphs had lower fat content than spring nymphs indicating that they are younger in age. All these results support the direct development hypothesis, and they contradict the delayed diapause hypothesis. In areas with a bimodal phenology, the DON and risk of tick-borne disease will increase 1 year later in the fall and 2 years later in the spring. Studies on the population ecology of *I. ricinus* should consider that the fall and peak that bookend the same winter contain the same generation of ticks. Studies on the annual abundance of *I. ricinus* should calculate this variable over the tick year (start of the fall in year *y*-1 to end of the summer of year *y*) and not the calendar year.

## Supporting information

Electronic supplemental material

## Additional files

**Additional file 1: Section 1.** Interpolation of the climate data from the weather stations. **Section 2.** Goodness of fit for the best model. **Section 3.** Effect of elevation site on the density of *I. ricinus* nymphs. **Section 4.** Correlation plots between the fall peak and the spring peak with different time lags. **Section 5.** Smoother function of calendar day predicts the bimodal phenology of *I. ricinus* nymphs at the four elevation sites. **Section 6.** Sequential modelling approach. **Section 7.** Comparison of the beech masting variables with different time lags and the inter-annual variation in the spring and fall peaks of the DON. **Section 8.** Full AIC-based model selection analysis. (DOCX).

## Abbreviations

AIC: Akaike information criterion
ASL: Above sea level
BM: Beech masting
DIN: Density of infected nymphs
DON: Density of nymphs
GAMs: Generalized additive models
PR: Precipitation
RH: Relative humidity
SD: Saturation deficit
SN: Snow fall
T: Temperature
WMO: World Meteorological Organization

## Acknowledgements

We would like to thank Lise Gern for her financial support and for generously giving us these data. This study was part of the PhD thesis of Cindy Bregnard.

## Declarations

### Ethics approval and consent to participate

Not applicable

### Consent for publication

Not applicable

### Availability of data and materials

The raw data for this study are stored in the Additional file 2. The climate data are available from the Climap-net database of the Federal Office for Meteorology and Climatology (http://www.meteosuisse.admin.ch/home/service-et-publications/conseil-et-service/portail-de-donnees-dedie-aux-specialistes.html). The MASTREE database is available in the Ecology – Ecological Society of America repository (http://onlinelibrary.wiley.com/doi/10.1002/ecy.1785/suppinfo).

## Funding

This study was supported by grants obtained by Lise Gern from the Swiss National Science Foundation: FN 32-57098.99, FN 3200B0-100657, FN 320000-113936 and FN 310030-127064 and by grants obtained by Lise Gern from the Federal Office of Public Health National Reference Center: 2009/10 (Projekt (911) 316) and 2011/13 (11.006911/ 304.0001-707). The doctoral salary of Cindy Bregnard was supported by the University of Neuchâtel. This research was also supported by two grants awarded to Maarten J. Voordouw, a Discovery Grant from the Natural Sciences and Engineering Research Council of Canada (RGPIN-2019-04483) and an Establishment Grant from the Saskatchewan Health Research Foundation (4583).

### Authors’ contributions

OR collected the ticks and the meteorological data in the field and managed the data. CH generated the data on the fat content in *I. ricinus* nymphs. CB and MJV analyzed the data and wrote the manuscript. OK provided critical insight into how diapause structures the bimodal phenology of *I. ricinus* in continental Europe. KB provided critical insight into modelling the DON as a function of climate variables averaged over different temporal windows and with different time lags. All authors read and approved the final version of the manuscript.

