## Supplementary material for "Beech tree masting explains the inter-annual variation in the fall and spring peaks of *Ixodes ricinus* ticks with different time lags": Electronic supplemental material

**ELECTRONIC SUPPLEMENTARY MATERIAL**

SECTION 1 – Interpolation of the climate data from the weather stations

**Methods:** We obtained climate data from the Federal Office of Meteorology and Climatology MeteoSwiss using the CLIMAP-net application. Climate data were obtained from two weather stations that are close to our four elevation sites and that are located at 485 m ASL in Neuchâtel and at 1136 m ASL in Chaumont. To create a climate profile that was specific for each of the four elevation sites, we interpolated the values between the two weather stations using the relative elevation distance of each elevation site to the two weather stations, as we have done previously (Bregnard et al. 2020, Bregnard et al. 2021). For example, the total elevation distance between the Neuchâtel and Chaumont weather stations is 651 meters, and the elevation distance between the top site and the Neuchâtel weather station is 620 meters, which represents 95.2% of the elevation distance. Thus, the climate at the top site is expected to be more similar to the Chaumont weather station (95.2%) compared to the Neuchâtel weather station (4.8%), whereas the reverse would be true for the low site. For each elevation site, the mean daily temperature, relative humidity, saturation deficit, precipitation and snowfall were calculated based on the interpolating percentages (Table S1).

Table S1. Interpolation of the climate data from the two weather stations. Shown are the site, elevation (in meters), elevation distance with the Neuchâtel weather station (Dist 1, in meters), elevation distance between Neuchâtel and Chaumont weather station (Dist 2, in meters), the interpolating percentage from the Neuchâtel weather station (Neuchâtel), and the interpolating percentage from the Chaumont weather station (Chaumont).

| **Site** | **Elevation** | **Dist 1** | **Dist 2** | **Neuchâtel** | **Chaumont** |
|---|---|---|---|---|---|
| Top | 1073 | 620 | 651 | 4.8% | 95.2% |
| High | 900 | 447 | 651 | 31.3% | 68.7% |
| Medium | 740 | 287 | 651 | 55.9% | 44.1% |
| Low | 620 | 167 | 651 | 74.3% | 25.7% |

SECTION 2 – Goodness of fit for the best model

**Methods:** We used the gam.check () function in R to assess the goodness of fit for the best GAM of the model selection table of the main manuscript (model 1 in Table 5 in the main manuscript).

**Results for nymphal abundance:** For the DON, the residuals of the best GAM (model 1 in Table 5 in the main manuscript) met the assumptions of the method (Figure S1) and therefore confirmed the goodness of fit.

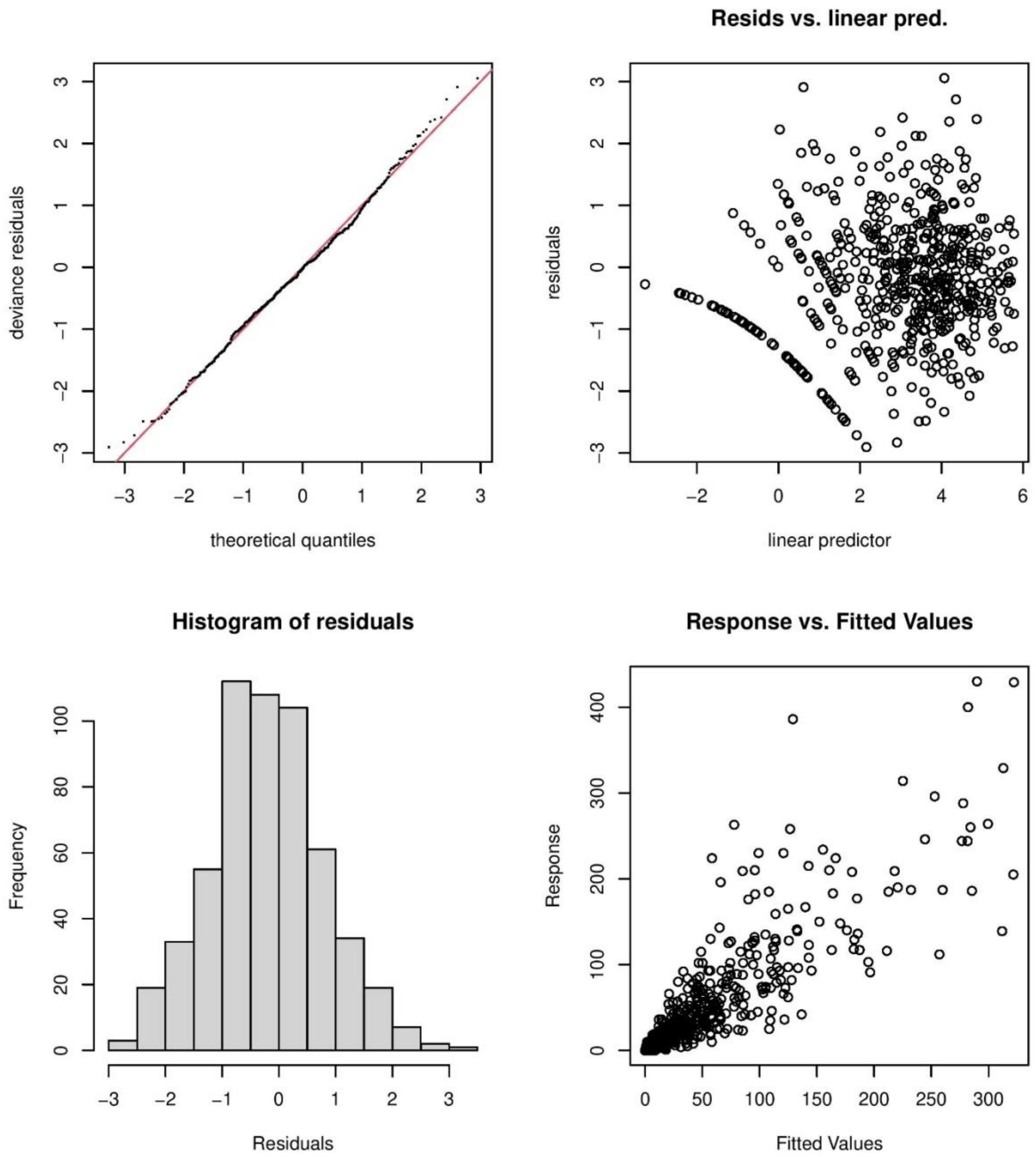

Figure S1. Goodness of fit assumptions for the best GAM in the model selection table of the main manuscript (model 1 in Table 5 in the main manuscript). The four plots generated by the gam.check() function confirmed that the residuals of the best model generally fit the assumptions of the GAM.

SECTION 3 – Effect of elevation site on the density of *I. ricinus* nymphs

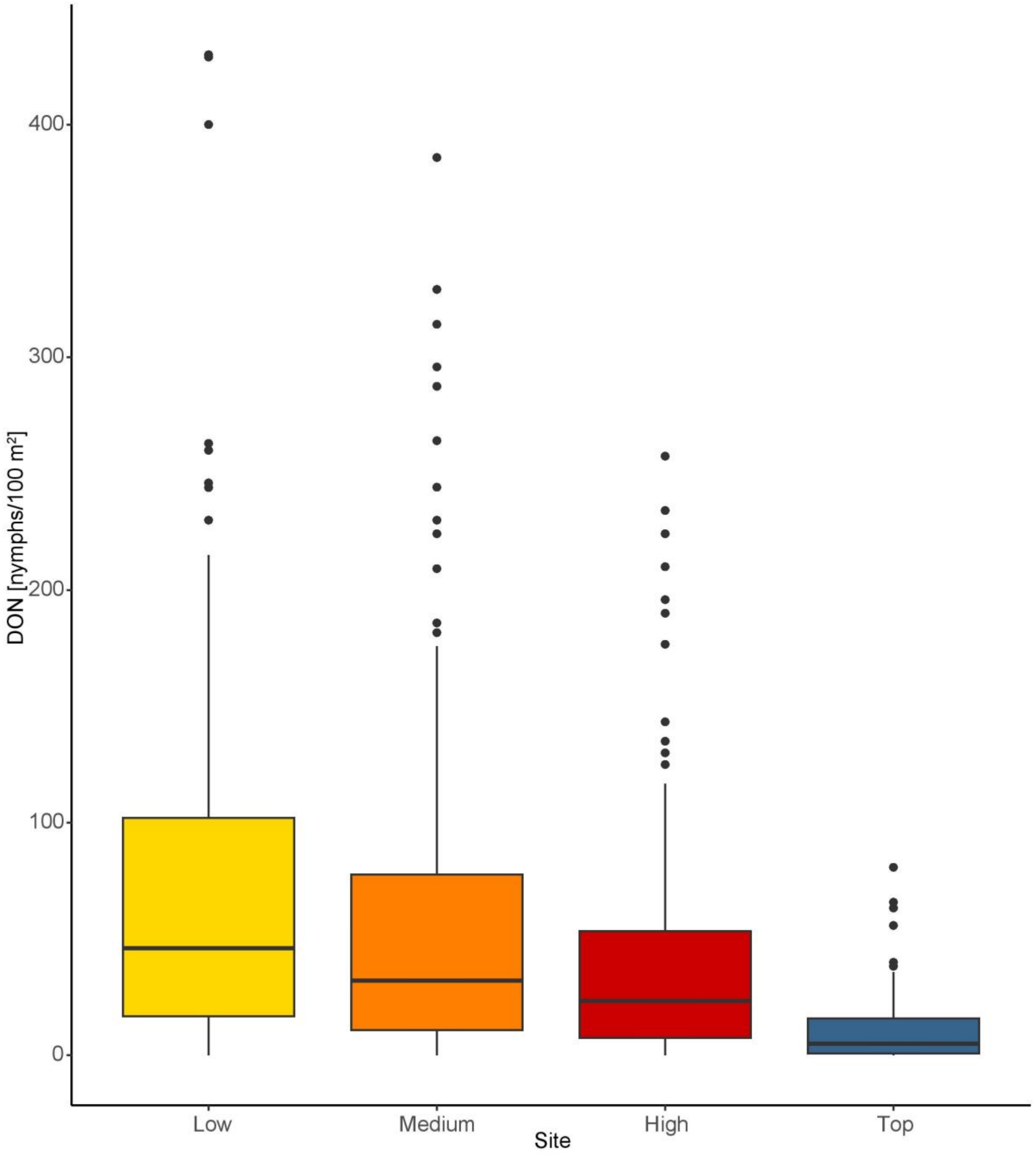

Figure S2. Effect of elevation on the density of nymphs (DON). The DON is an estimate of the number of questing *I. ricinus* nymphs per 100 $m^2$ sampled by the dragging method each month. The boxplot shows the medians (black line), the 25th and 75th percentiles (edges of the box), the minimum and maximum values (whiskers), and the outliers (solid circles).

SECTION 4 – Correlation plots between the fall peak and the spring peak with different time lags

**Methods:** Under the direct development hypothesis, the fall peak in year *y*-1 should be correlated with the spring peak in year *y*. In contrast, under the delayed diapause hypothesis, the fall peak in year *y* should be strongly correlated with the spring peak in year *y*. To compare these two competing hypotheses, we created scatter plots of the fall peak versus the spring peak with different time lags and calculated the Pearson correlation coefficient. As an additional control, we tested whether the fall peak in year y+1 was correlated with the spring peak in year y, even though this situation is not consistent with any particular hypothesis.

**Results:** The fall peak in year *y*-1 was strongly correlated with the spring peak in year *y* for the low and medium sites (Figure 5 in the main manuscript). In contrast, the fall peak and the spring peak in the same calendar year were not correlated (Figure S3). Similarly, the fall peak in year y+1 was not correlated with the spring peak in year y (Figure S4) and can be thought of as an additional negative control. Taken together, these correlation plots strongly provide strong evidence for the direct development hypothesis and that the tick year starts in the fall and ends the following summer. Most studies that analyze inter-annual variation in tick abundance calculate the total annual tick abundance over the calendar year. This approach is clearly wrong for our study location.

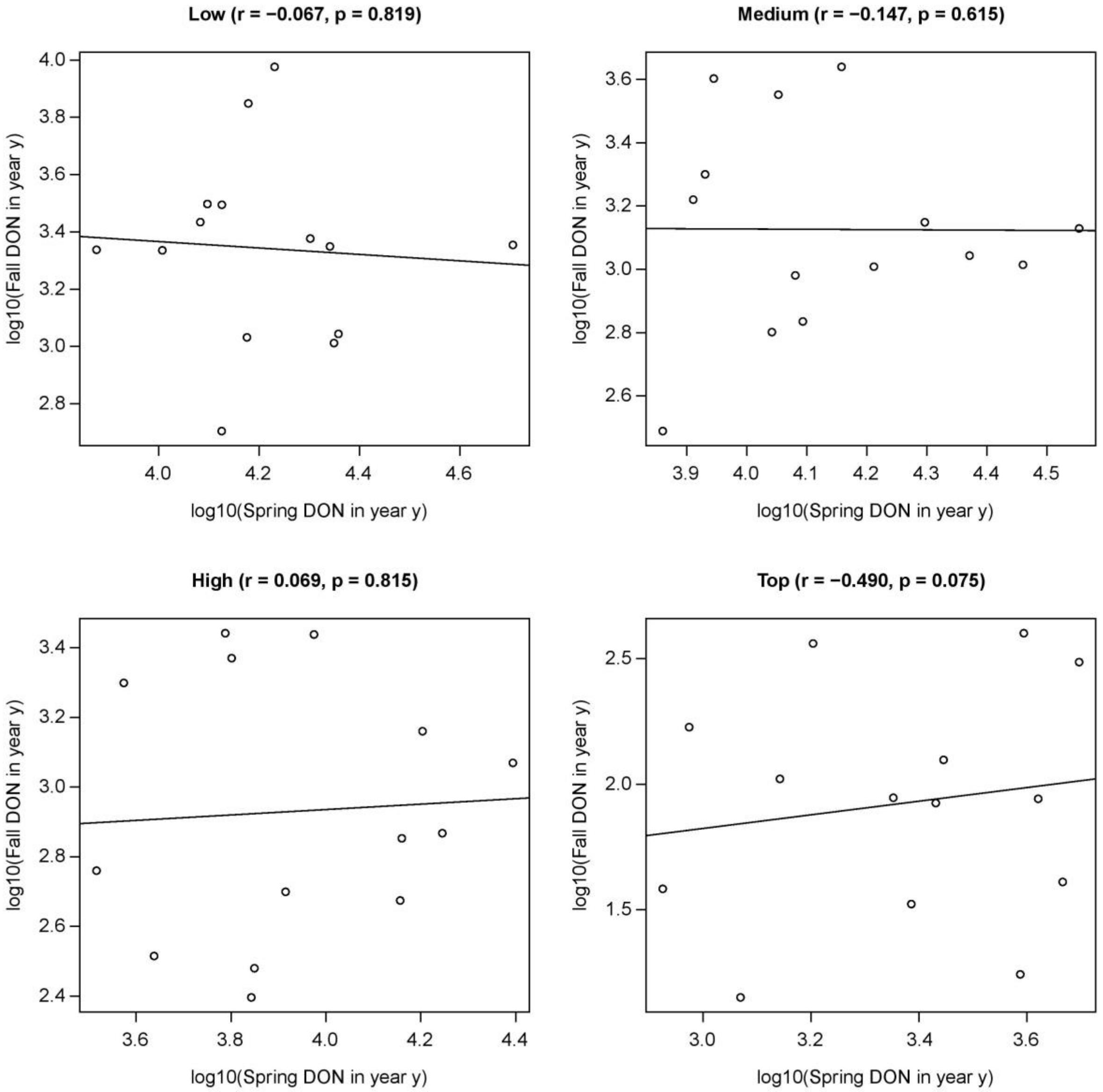

Figure S3. Correlation plot showing the relationship between the fall peak and the spring peak in the same calendar year for each of the four elevation sites. The fall peak in year *y* is not correlated with the spring peak in year *y*. The absence of significant correlations contradicts the delayed diapause hypothesis, which predicts that the fall peak in year *y* will be correlated with the spring peak in year *y*.

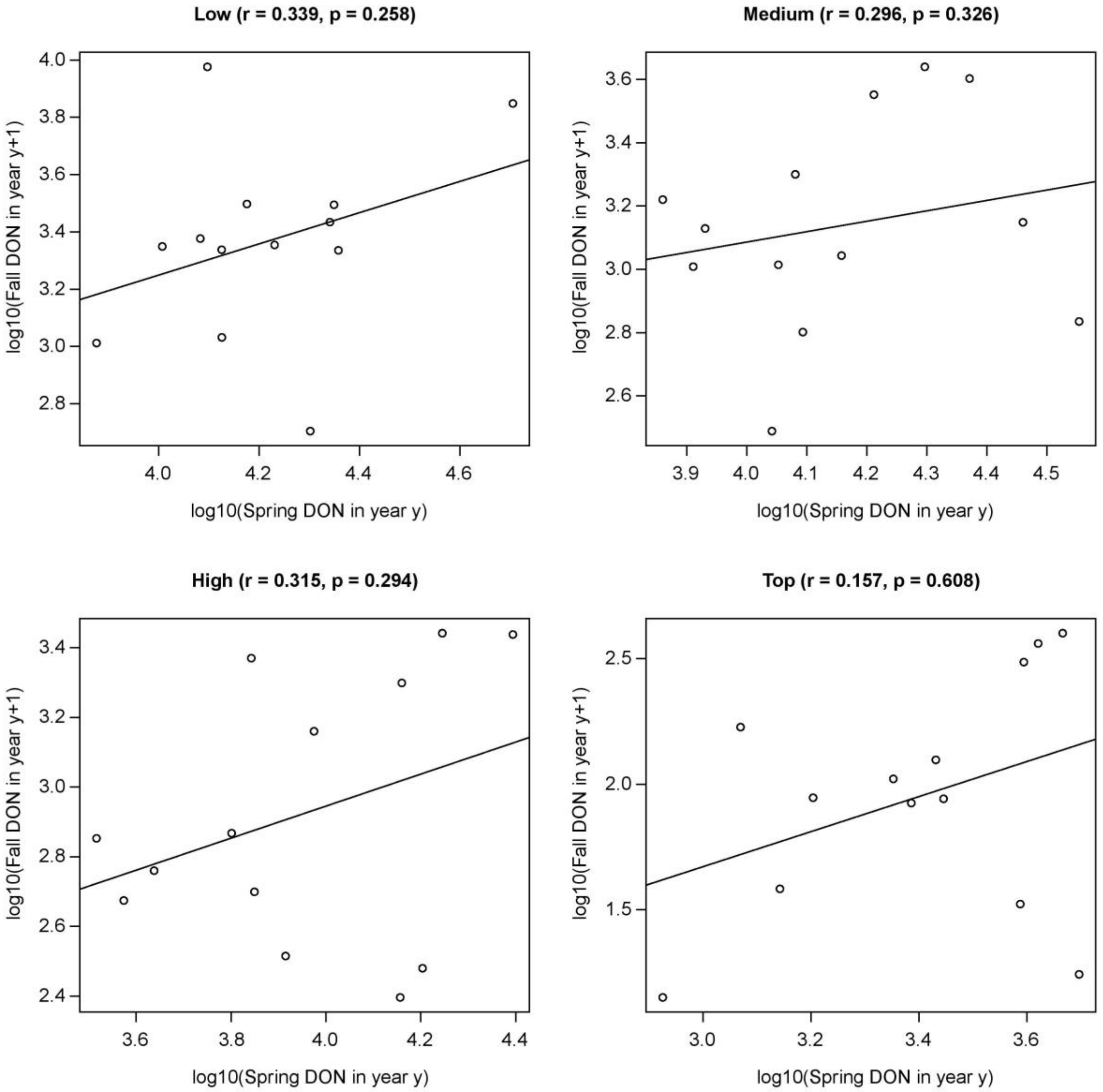

Figure S4. Correlation plot showing the relationship between the fall peak in year y+1 and the spring peak in year y for each of the four elevation sites. The fall peak in year y+1 is not correlated with the spring peak in year y. The correlation between the spring peak in year *y* and the fall peak in year *y*+1 is not predicted by any hypothesis about the diapause and phenology of *I. ricinus* nymphs. Figure S4 can be thought of as a control for Figure S3 and Figure 5 in the main manuscript.

SECTION 5 – Smoother function of calendar day predicts the bimodal phenology of *I. ricinus* nymphs at the four elevation sites.

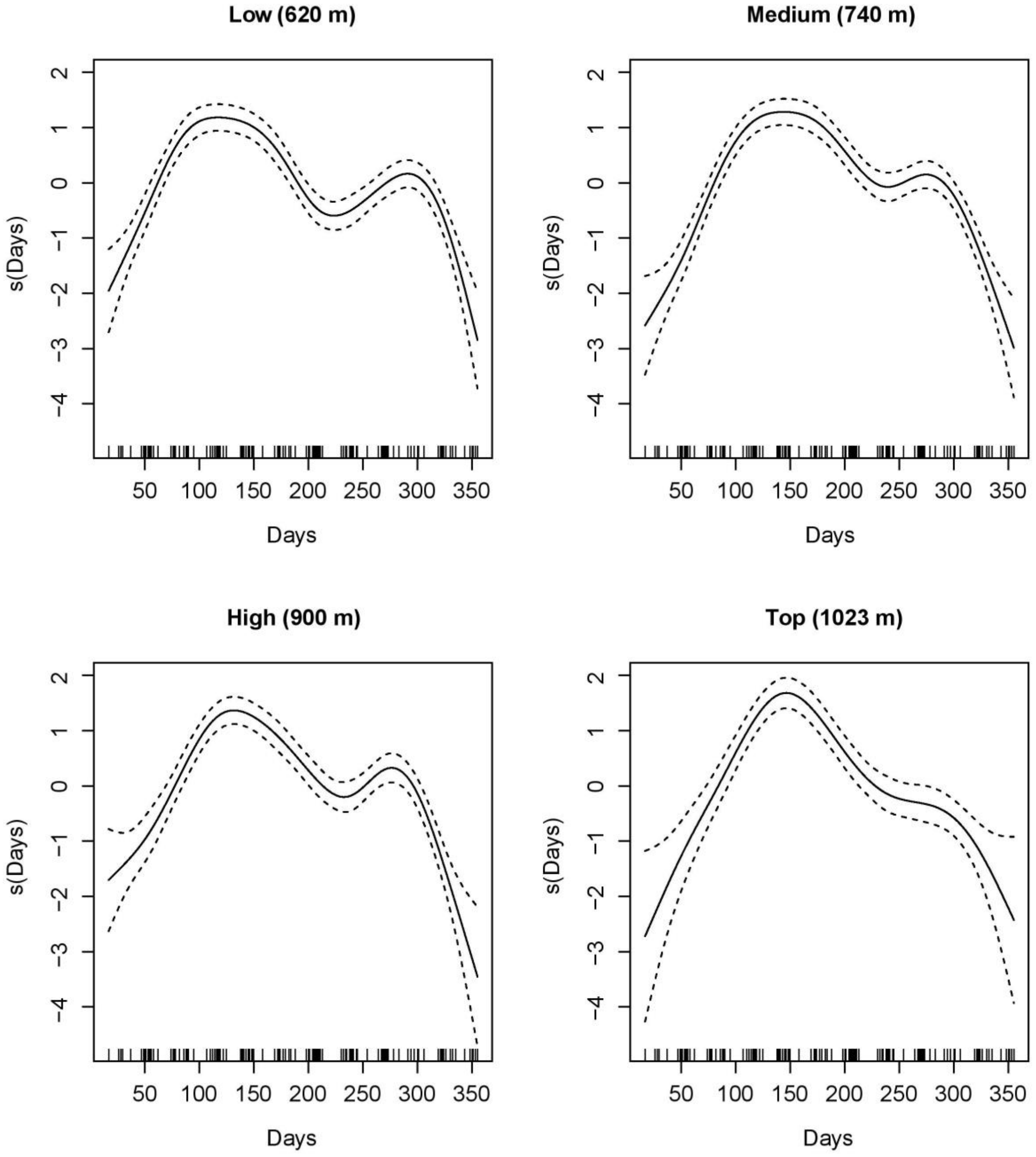

Figure S5. Visual interpretation of the smoother of the calendar day for each of the four elevation sites (partial $r^2$ = 32.1%). The smoother function of the calendar day recreates the bimodal phenology for the low, medium, and high elevation sites, whereas the top elevation site has a unimodal phenology.

SECTION 6 – Sequential modelling approach

To determine the best model for explaining variation in the monthly DON over the 14 years of the study (2004 to 2017) at the four elevation sites on Chaumont Mountain, we used a sequential modelling approach. We used our previous work on the same data set as starting point (Bregnard et al. 2020, Bregnard et al. 2021). This work had calculated the cumulative nymphal density (CND) for each calendar year (i.e., the CND values in Table 3 in the main manuscript) and found that the inter-annual variation in the CND was best explained by elevation site, year, site:year interaction, and the beech masting score 2 years prior. We therefore included these four explanatory variables in our set of starting models, as well as the previously mentioned smoother function of the calendar day to model the bimodal non-linear phenology of the DON (Figure S5). The original model set contained 23 models that were composed of different combinations of these 5 explanatory variables. We applied non-parametric smoother functions to all the explanatory variables (except site and site:year) and allowed for interactions with site. According to AIC-based model selection, the best model had an AIC score of 4422.1, 100% of the support, and an $r^2$ value of 76.7% (Table S2). As expected this best model (hereafter referred to as the base model 1) contained all 5 explanatory variables: site, smoothed function of year, site:year interaction, smoothed function of beech masting score, and smoothed function of calendar day. The relationship between the DON and year as well as the relationship between the DON and beech masting score both followed a linear relationship. We therefore removed the smoother functions for year and beech masting score, and all explanatory variables (except calendar day) were modelled as linear effects in base model 2. This base model 2 had an AIC score of 4482.1 and had an $r^2$ value of 68.3% (Table S2). The $r^2$ value of base model 1 (76.7%) is higher than that of base model 2 (68.3%) because non-parametric smoother functions of the explanatory variables always have better fit than the corresponding parametric functions. In summary, base model 2 confirmed our previous analyses (Bregnard et al. 2020, Bregnard et al. 2021) and that we could model the complex bimodal non-linear phenology of the monthly DON using a smoother function of calendar day.

Using base model 2 as a starting point, we wanted to test which of the 66 climate variables explained additional variation in the DON over the 14 years of the study. As shown in Table 2 in the main manuscript, the 66 climate variables included 3 field-measured climate variables on the day of tick sampling, 15 annual means from the weather stations (5 climate variables * 3 time lags), and 48 seasonal means from the weather stations (4 climate variables * 4 seasons * 3 time lags). We compared a set of 66 models; each model included the base 2 model and one of the 66 climate variables. We applied non-parametric smoothers to each of the 66 climate variables to account for non-linear effects. According to AIC-based model selection, the best model (hereafter referred to as base model 3) had an AIC score of 4389.8, 100% of the support, and an $r^2$ value of 69.5% (Table S2). Base model 3 contained the base model 2 and the field-measured temperature on the day of tick sampling.

The DON had a quadratic relationship with the field-measured temperature on the day of tick sampling that did not appear to differ between the four elevation sites. In base model 4, we therefore replaced the non-parametric smoother function of the field-collected temperature with a parametric quadratic function. Base model 4 had an AIC score of 4386.8 and had an $r^2$ value of 67.1% (Table S2). Again, the $r^2$ value of base model 3 (69.5%) is higher than that of base model 4 (67.1%) because the site-specific non-parametric smoother function of the field-measured temperature always has a better fit than the site-invariant quadratic function. Using base model 4 as a starting point, we tested whether any of the 63 annual or seasonal climate variables explained additional variation in the DON over the 14 years of the study. We compared a set of 63 models; each model included base model 4 and one of the 63 annual or seasonal climate variables. We again applied non-parametric smoothers to each of these 63 climate variables to account for non-linear effects. According to AIC-based model selection, the three best models (hereafter referred to as base models 5a, 5b, and 5c) had AIC scores of 4344.1, 4347.6 and 4353.6, supports of 85.0%, 14.0%, and 1.0%, and $r^2$ values of 68.2%, 72.7%, and 70.7%, respectively (Table S2). Models 5a, 5b, and 5c each contained base model 4 and the annual snow fall in year *y*-1 ($SN_{Y1}$), mean saturation deficit of the summer in year *y* ($SD_{S0}$) and mean seasonal relative humidity of the summer in year *y* ($RH_{S0}$), respectively. We created base models 6a, 6b, and 6c, by replacing the site-specific non-parametric smoother functions in base models 5a, 5b, and 5c with an interaction between elevation site and a quadratic function of the respective climate variable ($SN_{Y1}$, $SD_{S0}$, and $RH_{S0}$). Base models 6a, 6b, and 6c had AIC scores of 4352.3, 4344.1 and 4355.2, supports of 2.0%, 98.0%, and 0.0%, and $r^2$ values of 67.7%, 71.4%, and 69.6%, respectively (Table S2). Again, the $r^2$ values of base models 5a, 5b, and 5c (68.2%, 72.7%, and 70.7%) were higher than that of base models 6a, 6b, and 6c (67.7%, 71.4%, and 69.6%) for the previously mentioned reasons.

In the main manuscript, base models 6b, 6a, and 6c were renamed as models 1, 2, and 3 in Table 5 in the main manuscript. We ran a final set of 13 models to determine whether these three climate variables ($SN_{Y1}$, $SD_{S0}$, and $RH_{S0}$) were best modelled using linear or quadratic terms (models 1, 2, and 3 in Table 5 in the main manuscript). Interestingly, there was no support for any of the annual climate variables, which had been important in our previous work on the inter-annual variation of the DON and the DIN (Bregnard et al. 2020, Bregnard et al. 2021).

Table S2. Changes in the adjusted $r^2$ values for the best models during the sequential modelling approach. The explanatory variables were site (S), year (Y), site:year interaction (S:Y), calendar day, beech mast score with different time lags for spring and fall peak ($BM_{2/1}$), field-measured temperature on the day of tick sampling (t), Annual snowfall in the previous year ($SN_{Y1}$), mean seasonal SD of the summer in the present year ($SD_{S0}$), and mean seasonal RH of the summer in the present year ($RH_{S0}$). The explanatory variables were either modelled as non-parametric smoother functions or as parametric functions (e.g., linear effects or quadratic effects). To model the bimodal non-linear phenology of the DON, a site-specific smoother function was applied to the calendar day, s(day, by = S). Shown for each model are the explanatory variable compared between models (Var.), model identity (ID), model structure (see Table 2 in the main manuscript for the list of acronyms of the explanatory variables), and adjusted $r^2$ value ($r^2$).

| **Base** | **Type** | **Model structure** | **AIC** | **$r^2$ (%)** |
|---|---|---|---|---|
| 1 | Smoother | DON ~ [a] + S + s(Y, by = S) + s($BM_{2/1}$, by = S) | 4422.1 | 76.7 |
| 2 | Parametric | DON ~ [a] + S + Y + S:Y + $BM_{2/1}$ | 4482.1 | 68.3 |
| 3 | Smoother | DON ~ [b] + s(t, by = S) | 4389.8 | 69.5 |
| 4 | Parametric | DON ~ [b] + t + $t^2$ | 4386.8 | 67.1 |
| 5a | Smoother | DON ~ [c] + s($SN_{Y1}$, by = S) | 4344.1 | 68.3 |
| 6a | Parametric | DON ~ [c] + $SN_{Y1}$ + $SN_{Y1}^2$ + S:$SN_{Y1}$ + S: $SN_{Y1}^2$ | 4352.3 | 67.7 |
| 5b | Smoother | DON ~ [c] + s($SD_{S0}$, by = S) | 4347.6 | 72.7 |
| 6b | Parametric | DON ~ [c] + $SD_{S0}$ + $SD_{S0}^2$ + S:$SD_{S0}$ + S:$SD_{S0}^2$ | 4344.1 | 71.4 |
| 5c | Smoother | DON ~ [c] + s($RH_{S0}$, by = S) | 4353.6 | 70.7 |
| 6c | Parametric | DON ~ [c] + $RH_{S0}$ + $RH_{S0}^2$ + S:$RH_{S0}$ + S:$RH_{S0}^2$ | 4355.2 | 69.6 |

[a] All models contain the smoother function of calendar day: s(day, by = S)

[b] All models contain the following terms: s(day, by = S)+S+Y+S:Y+$BM_{2/1}$

[c] All models contain the following terms: s(day, by = S)+S+Y+S:Y+$BM_{2/1}$+t+$t^2$

SECTION 7 – Comparison of the beech masting variables with different time lags and the inter-annual variation in the spring and fall peaks of the DON

**Methods:** We created three different explanatory variables, $BM_{2/2}$, $BM_{1/1}$, and $BM_{2/1}$, for the beech masting (BM) score. $BM_{2/2}$ assumes there is a 2-year time lag between BM and the spring and fall peaks of nymphs. $BM_{1/1}$ assumes there is a 1-year time lag between BM and the spring and fall peaks of nymphs. $BM_{2/1}$ assumes there is a 2-year time lag and a 1-year time lag between BM and the spring and fall peak, respectively. $BM_{2/2}$ is consistent with the developmental diapause hypothesis, $BM_{2/1}$ is consistent with the direct development hypothesis, and $BM_{1/1}$ is not consistent with either hypothesis. To determine which of these three beech masting variables best explained the inter-annual variation in the spring and fall nymphal peaks, we re-ran the best model in the main manuscript (model 1 of Table 5 in the main manuscript), and included either $BM_{1/1}$, $BM_{2/2}$, or $BM_{2/1}$ as the explanatory variable for beech masting. As a side note, we conducted a similar comparison at the beginning of our sequential modelling approach (e.g., 639 different models), which we present in the results of this section as well.

**Results:** The best model had a support of 100.0%, explained 71.4% of the variation in the DON, and contained the masting explanatory variable $BM_{2/1}$ (Table S3). The two models containing $BM_{1/1}$ and $BM_{2/2}$ each had a support of 0.0%, explained 60.3% and 64.6% of the variation in the DON, and the model AIC value was 166.3 and 137.2 units higher, respectively (Table S3). When we conducted a similar analysis at the beginning of our sequential modelling approach (e.g., 639 different models), we found the same result. The best model had a support of 100.0%, explained 72.4% of the variation in the DON, and contained $BM_{2/1}$ (Table S3). The two models containing $BM_{1/1}$ and $BM_{2/2}$ each had a support of 0.0%, explained 66.6% and 76.2% of the variation in the DON, and the model AIC value was 175.0 and 84.0 units higher, respectively (Table S3). Based on these model comparisons, we are confident that the beech masting index with a 2-year time lag for the spring peak and a 1-year time lag for the fall peak best explained the inter-annual variation in the DON.

Table S3. Statistical analysis to test whether different time lags explain the inter-annual variation in the spring and fall peaks of *I. ricinus* nymphs. Model selection results are shown for the generalized additive model (GAM) with negative binomial errors of the monthly density of nymphs (DON). This analysis compared three different beech masting (BM) variables: $BM_{2/2}$, $BM_{1/1}$, and $BM_{2/1}$; the subscripts 'i/j' refer to the time lag (in years) for the spring nymphal peak and the fall nymphal peak, respectively. We compared these three BM variables on a common background model at the beginning and end of our sequential modelling approach. At the beginning, we used background model b, which contained site and the site-specific smoother function of the calendar day. At the end, we used background model a, which was the best model in the main text (model 1 in Table 5 in the main manuscript). In both analyses, the model with $BM_{2/1}$ had all of the support, whereas the models with $BM_{2/2}$ and $BM_{1/1}$ had no support. The models are ranked according to their Akaike Information Criterion (AIC). Shown for each model are the model rank (Rank), model structure (see below for explanation of explanatory variables), model degrees of freedom (Df), log-likelihood (logLik), Akaike information criterion (AIC), difference in the AIC value from the top model (ΔAIC), model weight (Weight1), cumulative weight (Weight2), and adjusted r-squared value ($r^2$).

| Rank | Model structure | Df | logLik | AIC | ΔAIC | Weight1 | Weight2 | $r^2$ |
|---|---|---|---|---|---|---|---|---|
| 1 | …[a] + $BM_{2/1}$ | 49 | -2117.8 | 4344.1 | 0.0 | 100.0 | 100.0 | 71.4 |
| 2 | …[a] + $BM_{2/2}$ | 49 | -2186.4 | 4481.3 | 137.2 | 0.0 | 100.0 | 64.6 |
| 3 | …[a] + $BM_{1/1}$ | 49 | -2201.3 | 4510.4 | 166.3 | 0.0 | 100.0 | 60.3 |
| 1 | …[b] + $BM_{2/1}$ | 47 | -2204.3 | 4512.7 | 0.0 | 100.0 | 100.0 | 72.4 |
| 2 | …[b] + $BM_{2/2}$ | 48 | -2245.5 | 4596.7 | 84.0 | 0.0 | 100.0 | 76.2 |
| 3 | …[b] + $BM_{1/1}$ | 40 | -2299.6 | 4687.7 | 175.0 | 0.0 | 100.0 | 66.6 |

[a] The model structure of explanatory variables was $S+Y+S:Y+t+t^2+s(day, by = S)+SDS0+SD_{S0}^2+S:SD_{S0}+S:SD_{S0}^2$.
[b] The model structure of explanatory variables was S+s(day, by = S).

SECTION 8 – Full AIC-based model selection analysis

**Methods:** To identify the best model, we used a model selection approach based on the Akaike information criterion (AIC). Models were ranked according to their AIC values and the Akaike weights, which indicate the percent support, were calculated for each model. We used the Akaike weights to calculate the model-averaged parameter estimates and their 95% confidence intervals (CIs).

**Results:**

**Full model selection table:** The full model selection table for all 13 models is presented in Table S4. For the monthly DON, the best model had a support of 98.0%, and explained 71.4% of the variation in the DON. This model contained the explanatory variables of site, year, site:year interaction, beech mast score with different time lags for spring and fall peak ($BM_{2/1}$), quadratic function of the temperature on the day of tick sampling (t and $t^2$), quadratic function of the weather station mean seasonal SD of the summer in the present year ($SD_{S0}$, $SD_{S0}^2$, site:$SD_{S0}$, and site:$SD_{S0}^2$), and the site-specific smoother function of the calendar day (Table S4). The second-best and third-best model had 2% and 0% of the support respectively. The second-best and third-best model were identical to the top model except that it contained the annual snow fall ($SN_{Y1}$) in the previous year and the mean seasonal RH of the summer in the present year ($RH_{S0}$) instead of $SD_{S0}$.

**Support for the individual explanatory variables:** The supports for the 11 most important individual explanatory variables are shown in Table S5 and were as follows: site (100.0%), year (100.0%), site:year interaction (100.0%), beech mast score with different time lags for spring and fall peak ($BM_{2/1}$; 100.0%), temperature on the day of tick sampling (support is 100.0% for both t and $t^2$), smoothed function of the calendar day (100.0%), weather station mean seasonal SD of the summer in the present year (support is 97.8% – 97.9% for $SD_{S0}$, $SD_{S0}^2$, site:$SD_{S0}$ interaction, and the site:$SD_{S0}^2$ interaction). None of the other explanatory variables had a support > 2.0% (Table S5).

**Model-averaged parameter estimates:** To determine the effect of the explanatory variables on the DON, we present the model-averaged parameter estimates on the log scale (and their 95% confidence intervals; Table S6). We also back calculated the effect sizes of the explanatory variables on the DON on the original scale with respect to the following reference conditions: the site was low elevation, the year was 2004, and the beech tree mast index ($BM_{2/1}$) was given a value of 1. The model-averaged parameter estimates (Table S6) are virtually identical to the parameter estimates of the top model (Table 7 in the main manuscript) because the top model has 98.0% of the weight.

The interaction between elevation site and year indicated that the change in the DON over time differed between the four elevation sites (Table S6). Over the 14-year period (2004 – 2017), the DON increased at the low elevation site (slope = 0.056 per year, 95% CI = 0.029 to 0.083) and the medium elevation site (Medium – Low contrast of the slope = -0.034, 95% CI = -0.074 to 0.005), but the DON decreased at the high elevation site (High – Low contrast of the slope = -0.091, 95% CI = -0.128 to -0.053) and the top elevation site (Top – Low contrast of the slope = -0.142, 95% CI = -0.185 to -0.099). Over the 14-year period (2004 – 2017), the DON increased by 119.0% and 36.1% at the low and medium elevation sites but decreased by 38.7% and 70.0% at the high and top elevation sites, respectively.

The beech mast score with different time lags (2 years versus 1 year) for the spring and fall peak ($BM_{2/1}$) had a positive and significant effect on the DON (slope = 0.232 per class, 95% CI = 0.199 to 0.264; Table S6). Increasing the beech mast score from 1 (poor mast) to 5 (full mast) increased the DON by 152.9% at each of the four elevation sites on Chaumont Mountain.

The slope of the linear effect of the field-measured temperature (i.e., the temperature measured on the day of tick sampling) on the DON was positive and significant (slope = 9.608 per °C, 95% CI = 7.189 to 12.028; Table S6), indicating that the DON increased with temperature over the range of observed values (-5°C to 30°C). The slope of the quadratic effect of the field-measured temperature on the DON was negative and significant (slope = -8.828, 95% CI = -10.652 to -7.004; Table S6) indicating that the DON plateaued once the field-measured temperature reached ~ 30°C.

The $SD_{S0}$ is the mean SD during the summer (1 June to 31 August) of the same year as the DON (i.e., no time lag). Thus, the $SD_{S0}$ represents the saturation deficit during the last 3 months of the spring nymphal peak, just before the start of the fall peak. The significant interaction between elevation site and $SD_{S0}$ indicates that the relationship between the DON and the $SD_{S0}$ differed between the four elevation sites (Table S6). The relationship between the DON and the $SD_{S0}$ was positive linear at the low and high elevation sites, and negative quadratic at the medium and top elevation sites. At the medium and top elevation sites, the DON decreased once the $SD_{S0}$ reached ~ 5.5 mmHg and ~ 4.0 mmHg, respectively.

In summary, the DON had a bimodal phenology at the low, medium, and high elevation sites and a unimodal phenology at the top elevation site. The DON increased over time at the low and medium elevation sites, whereas it decreased at the high and top elevation sites. The DON increased significantly with beech masting index and importantly, the time lag differed between the two nymphal peaks with a 2-year time lag for the spring peak and a 1-year time lag for the fall peak. This result provides strong evidence that the spring and fall nymphal peaks are distinct cohorts that are born in different years, which supports the direct development hypothesis and not the developmental diapause hypothesis. The DON increased with the field-measured temperature but reached a plateau at 30°C. Finally, the relationship between the DON and the seasonal SD of the summer in year y ($SD_{S0}$) was positive linear at the low and high elevation sites, and negative quadratic at the medium and top elevation sites.

Table S4. Full model selection results are shown for the generalized additive model (GAM) with negative binomial errors of the density of *I. ricinus* nymphs (DON) at the four elevation sites on Chaumont Mountain over 14 years (2004 to 2017). The explanatory variables were site, year, site:year interaction, day, beech mast score 2 years prior and 1 year prior for the spring and fall peak, respectively, field-measured temperature on the day of tick sampling, and 4 important weather station climate variables (see Overview of the preliminary analyses). All climate variables were coded as linear and quadratic fixed effects. A smoother function was applied to the covariate of calendar day. The models are ranked according to their Akaike Information Criterion (AIC). Shown for each model are the model rank (Rank), model structure (see below for explanation of explanatory variables), model degrees of freedom (Df), log-likelihood (logLik), Akaike information criterion (AIC), difference in the AIC value from the top model (ΔAIC), model weight (Weight1), cumulative weight (Weight2), and adjusted r-squared value ($r^2$).

| Rank | Model structure | Df | logLik | AIC | ΔAIC | Weight1 | Weight2 | $r^2$ |
|---|---|---|---|---|---|---|---|---|
| 1 | …[a]+$SD_{S0}$+$SD_{S0}^2$+S:$SD_{S0}$+S:$SD_{S0}^2$ | 49 | -2117.8 | 4344.1 | 0.0 | 98.0 | 98.0 | 71.4 |
| 2 | …[a]+$SN_{Y1}$+$SN_{Y1}^2$+S:$SN_{Y1}$+S:$SN_{Y1}^2$ | 49 | -2122.3 | 4352.3 | 8.2 | 2.0 | 100.0 | 67.7 |
| 3 | …[a]+$RH_{S0}$+$RH_{S0}^2$+S:$RH_{S0}$+S:$RH_{S0}^2$ | 49 | -2123.4 | 4355.2 | 11.0 | 0.0 | 100.0 | 69.6 |
| 4 | …[a]+$SD_{S0}$+S:$SD_{S0}$ | 45 | -2128.8 | 4356.7 | 12.6 | 0.0 | 100.0 | 69.6 |
| 5 | …[a]+$RH_{S0}$+S:$RH_{S0}$ | 45 | -2130.4 | 4359.7 | 15.6 | 0.0 | 100.0 | 67.7 |
| 6 | …[a]+$SD_{S0}$+$SD_{S0}^2$ | 43 | -2138.7 | 4371.4 | 27.3 | 0.0 | 100.0 | 69.2 |
| 7 | …[a]+$RH_{S0}$ | 42 | -2140.8 | 4373.2 | 29.1 | 0.0 | 100.0 | 70.0 |
| 8 | …[a]+$SD_{S0}$ | 42 | -2141.0 | 4373.6 | 29.5 | 0.0 | 100.0 | 70.6 |
| 9 | …[a]+$RH_{S0}$+$RH_{S0}^2$ | 43 | -2140.6 | 4374.9 | 30.7 | 0.0 | 100.0 | 69.3 |
| 10 | …[a]+$SN_{Y1}$+S:$SN_{Y1}$ | 44 | -2142.5 | 4383.0 | 38.8 | 0.0 | 100.0 | 67.4 |
| 11 | …[a]+$SN_{Y1}$ | 42 | -2147.7 | 4386.4 | 42.3 | 0.0 | 100.0 | 66.8 |
| 12 | …[a] | 41 | -2149.0 | 4386.8 | 42.6 | 0.0 | 100.0 | 67.1 |
| 13 | …[a]+$SN_{Y1}$+$SN_{Y1}^2$ | 42 | -2147.7 | 4388.8 | 44.7 | 0.0 | 100.0 | 66.7 |

[a] The model basic structure of explanatory variables were S+Y+S:Y+$BM_{2/1}$+t+$t^2$+s(day, by = S).

Table S5. The support for each of the 21 individual explanatory variables is shown for the GAMs of the DON. This support is calculated as the sum of the Akaike weights for all the models in the set that include that particular explanatory variable.

| Rank | Explanatory variable of interest | Support (%) |
|---|---|---|
| 1 | Site | 100.0 |
| 2 | Year | 100.0 |
| 3 | Site:Year | 100.0 |
| 4 | $BM_{2/1}$ | 100.0 |
| 5 | t | 100.0 |
| 6 | $t^2$ | 100.0 |
| 7 | s(Day, by = Site) | 100.0 |
| 8 | $SD_{S0}$ | 97.9 |
| 9 | $SD_{S0}^2$ | 97.8 |
| 10 | $S{:}SD_{S0}$ | 97.9 |
| 11 | $S{:}SD_{S0}^2$ | 97.8 |
| 12 | $SN_{Y1}$ | < 1.0 |
| 13 | $SN_{Y1}^2$ | < 1.0 |
| 14 | $RH_{S0}$ | < 1.0 |
| 15 | $RH_{S0}^2$ | < 1.0 |
| 18 | $S{:}SN_{Y1}$ | < 1.0 |
| 19 | $S{:}SN_{Y1}^2$ | < 1.0 |
| 20 | $S{:}RH_{S0}$ | < 1.0 |
| 21 | $S{:}RH_{S0}^2$ | < 1.0 |

Table S6. Model-averaged parameter estimates are shown for the GAMs with negative binomial errors of the DON. Shown are the parameter types, the parameter names, the parameter estimates, and the 95% confidence limits (LL = lower limit and UL = upper limit). Estimate 1 is averaged over all the models in the set. Estimate 2 is averaged over the subset of models with a cumulative support of 95%. The 95% confidence limits are for estimate 2.

| Type | Name | Estimate 1 | Estimate 2 | 95% LL | 95% UL |
|---|---|---|---|---|---|
| **Intercept** | **Low site** | **2.765** | **2.765** | **2.436** | **3.093** |
| **Contrast 1** | **Medium site** | **-0.496** | **-0.496** | **-0.878** | **-0.113** |
| Contrast 2 | High site | -0.149 | -0.149 | -0.655 | 0.357 |
| **Contrast 3** | **Top site** | **-1.894** | **-1.894** | **-2.674** | **-1.114** |
| **Slope 1** | **Year** | **0.056** | **0.056** | **0.029** | **0.083** |
| **Slope 2** | **$BM_{2/1}$** | **0.232** | **0.232** | **0.199** | **0.264** |
| **Slope 3** | **t** | **9.608** | **9.608** | **7.189** | **12.028** |
| **Slope 4** | **$t^2$** | **-8.828** | **-8.828** | **-10.652** | **-7.004** |
| Slope 5 | $SD_{S0}$ | -1.890 | -1.930 | -7.492 | 3.633 |
| Slope 6 | $SD_{S0}^2$ | 2.892 | 2.959 | -0.743 | 6.660 |
| Slope 7 | $SN_{Y1}$ | -0.352 | -21.692 | -52.189 | 8.804 |
| Slope 8 | $SN_{Y1}^2$ | -0.138 | -8.525 | -23.906 | 6.857 |
| Slope 9 | $RH_{S0}$ | 0.013 | 2.885 | -2.853 | 8.622 |
| Slope 10 | $RH_{S0}^2$ | 0.015 | 3.785 | 0.247 | 7.322 |
| Contrast 4 | Medium site:Year | -0.034 | -0.034 | -0.074 | 0.005 |
| **Contrast 5** | **High site:Year** | **-0.091** | **-0.091** | **-0.128** | **-0.053** |
| **Contrast 6** | **Top site:Year** | **-0.142** | **-0.142** | **-0.185** | **-0.099** |
| **Contrast 7** | **Medium site:$SD_{S0}$** | **10.859** | **11.088** | **3.837** | **18.339** |
| Contrast 8 | High site:$SD_{S0}$ | 11.453 | 11.694 | -4.210 | 27.598 |
| **Contrast 9** | **Top site:$SD_{S0}$** | **-22.966** | **-23.450** | **-43.662** | **-3.238** |
| **Contrast 10** | **Medium site:$SD_{S0}^2$** | **-14.676** | **-15.013** | **-22.600** | **-7.425** |
| Contrast 11 | High site:$SD_{S0}^2$ | -10.329 | -10.566 | -28.823 | 7.692 |
| **Contrast 12** | **Top site:$SD_{S0}^2$** | **-18.642** | **-19.070** | **-32.112** | **-6.029** |
| Contrast 13 | Medium site:$SN_{Y1}$ | 0.126 | 7.771 | -24.452 | 39.994 |
| Contrast 14 | High site:$SN_{Y1}$ | 0.336 | 20.702 | -9.942 | 51.345 |
| Contrast 15 | Top site:$SN_{Y1}$ | 0.409 | 25.233 | -5.712 | 56.177 |
| Contrast 16 | Medium site:$SN_{Y1}^2$ | -0.127 | -7.818 | -25.709 | 10.072 |
| Contrast 17 | High site:$SN_{Y1}^2$ | -0.102 | -6.313 | -22.481 | 9.856 |
| Contrast 18 | Top site:$SN_{Y1}^2$ | 0.143 | 8.785 | -6.969 | 24.539 |
| Contrast 19 | Medium site: $RH_{S0}$ | -0.052 | -11.872 | -19.761 | -3.982 |
| Contrast 20 | High site: $RH_{S0}$ | -0.066 | -14.987 | -26.128 | -3.846 |
| Contrast 21 | Top site: $RH_{S0}$ | 0.041 | 9.308 | -3.989 | 22.605 |
| Contrast 22 | Medium site:$RH_{S0}^2$ | -0.041 | -10.166 | -17.398 | -2.935 |
| Contrast 23 | High site:$RH_{S0}^2$ | -0.018 | -4.463 | -18.465 | 9.539 |
| Contrast 24 | Top site:$RH_{S0}^2$ | -0.053 | -13.297 | -21.843 | -4.750 |

| | | | | | |
|---|---|---|---|---|---|
| Contrast 25 | s(Day, low 1) | 0.537 | 0.537 | -0.619 | 1.692 |
| Contrast 26 | s(Day, low 2) | -8.227 | -8.227 | -11.195 | -5.258 |
| Contrast 27 | s(Day, low 3) | -2.464 | -2.464 | -3.258 | -1.669 |
| Contrast 28 | s(Day, low 4) | 4.359 | 4.359 | 2.209 | 6.510 |
| Contrast 29 | s(Day, low 5) | 1.518 | 1.518 | 0.598 | 2.437 |
| Contrast 30 | s(Day, low 6) | 3.358 | 3.358 | 1.449 | 5.266 |
| Contrast 31 | s(Day, low 7) | 1.090 | 1.090 | 0.523 | 1.658 |
| Contrast 32 | s(Day, low 8) | 12.083 | 12.083 | 7.203 | 16.963 |
| Contrast 33 | s(Day, low 9) | -1.737 | -1.737 | -3.746 | 0.272 |
| Contrast 34 | s(Day, medium 1) | 0.093 | 0.093 | -1.079 | 1.265 |
| Contrast 35 | s(Day, medium 2) | -3.908 | -3.908 | -6.942 | -0.874 |
| Contrast 36 | s(Day, medium 3) | -1.806 | -1.806 | -2.597 | -1.014 |
| Contrast 37 | s(Day, medium 4) | 2.055 | 2.055 | -0.122 | 4.231 |
| Contrast 38 | s(Day, medium 5) | 0.871 | 0.871 | -0.019 | 1.761 |
| Contrast 39 | s(Day, medium 6) | 1.189 | 1.189 | -0.699 | 3.076 |
| Contrast 40 | s(Day, medium 7) | 0.417 | 0.417 | -0.116 | 0.950 |
| Contrast 41 | s(Day, medium 8) | 6.997 | 6.997 | 2.008 | 11.986 |
| Contrast 42 | s(Day, medium 9) | -1.136 | -1.136 | -3.195 | 0.923 |
| Contrast 43 | s(Day, high 1) | 0.751 | 0.751 | -0.805 | 2.307 |
| Contrast 44 | s(Day, high 2) | -4.324 | -4.324 | -8.055 | -0.593 |
| Contrast 45 | s(Day, high 3) | -2.304 | -2.304 | -3.319 | -1.289 |
| Contrast 46 | s(Day, high 4) | 1.731 | 1.731 | -0.939 | 4.401 |
| Contrast 47 | s(Day, high 5) | 0.830 | 0.830 | -0.284 | 1.945 |
| Contrast 48 | s(Day, high 6) | 1.205 | 1.205 | -1.095 | 3.506 |
| Contrast 49 | s(Day, high 7) | 0.255 | 0.255 | -0.384 | 0.893 |
| Contrast 50 | s(Day, high 8) | 6.753 | 6.753 | 0.655 | 12.852 |
| Contrast 51 | s(Day, high 9) | -2.116 | -2.116 | -4.675 | 0.443 |
| Contrast 52 | s(Day, top 1) | -0.923 | -0.923 | -2.698 | 0.851 |
| Contrast 53 | s(Day, top 2) | -3.220 | -3.220 | -7.442 | 1.003 |
| Contrast 54 | s(Day, top 3) | -1.709 | -1.709 | -2.805 | -0.613 |
| Contrast 55 | s(Day, top 4) | 2.065 | 2.065 | -0.828 | 4.957 |
| Contrast 56 | s(Day, top 5) | 0.593 | 0.593 | -0.514 | 1.700 |
| Contrast 57 | s(Day, top 6) | 1.888 | 1.888 | -0.483 | 4.259 |
| Contrast 58 | s(Day, top 7) | 0.311 | 0.311 | -0.303 | 0.925 |
| Contrast 59 | s(Day, top 8) | 7.137 | 7.137 | 0.293 | 13.982 |
| Contrast 60 | s(Day, top 9) | 0.138 | 0.138 | -2.742 | 3.018 |
